# Giant spider neurons uncover a myelin-derived waste-internalizing canal system that fails in neurodegeneration

**DOI:** 10.64898/2026.02.17.706423

**Authors:** Ruth Fabian-Fine, Sondre M Brännare-Gran, Maya N Clough, Tyler A DeBoer, Isabella R DelVecchio, Matthew D Devers, Julie A Dragon, Heather E Driscoll, Zavier F Fuller, Keely M Hendricks, Alexander M Huesgen, Isabella C Joly, Landon G Kinal, Luwago K Kipingi, Kyleena J Lathram, Lydia A Kragh, James A Lubkowitz, Juliette S Majeske, Samantha C McDonough, Bree A McKinley, Oliver A Miatke, Jordan A Munro, Chloe M Paul, Oscar V Preisler, Calum J Reding, Kaylee C Scott, Ethan S St. Pierre, Hannah A Steen, Joshua E Syverson, Madeline E Waldt, Abigail E Whitley, Amy L Winchell, Adam L Weaver, Emily E Curd

## Abstract

The underlying causes for Alzheimer disease are presumed to lie in failed waste removal from the brain. However, the mechanisms by which waste is cleared from neurons, and how this system fails in neurodegeneration are poorly understood. A novel ‘glial-canal-hypothesis’ postulates that myelin-forming macroglia give rise to waste internalizing canals that project into neuronal somata and remove cellular debris in an aquaporin4-dependent manner. We postulate that abnormal swelling of the aquaporin4-expressing glial cells leads to spongiform abnormalities, gradual depletion, and death of associated neurons. Due to the novelty of this postulation little is known about the cellular architecture of this canal system that was first discovered in giant neurons of the wandering spider *Cupiennius salei*. Here we have utilized histological, ultrastructural and immunohistochemical methods to describe the structural foundation of this glial canal system in giant spider neurons in which waste-internalizing canals and associated structures are clearly visible. Sequencing the spider genome, we show compelling homologies of key proteins that are implicated in neurodegeneration between phylogenetically distant species. Based on this work, we provide a testable functional hypothesis regarding waste removal from neuronal somata and how this system fails in neurodegeneration. We highlight structural similarities of this system in rodent and human brain. Supported by the findings presented here we postulate that (*i*) neurodegeneration in *C. salei* may be caused by hypertrophic swelling of myelin-forming waste-internalizing macroglia, and (*ii*) that a similar canal system, although structurally modified, is likely highly conserved in the mammalian brain.

## INTRODUCTION

One of the most important topics in biomedical research today relates to the underlying mechanisms that trigger Alzheimer disease (AD) [1]. Based on the prevalent amyloid hypothesis [2], the underlying causes for AD lie in abnormal deposits of Amyloid β (Aβ) and tau protein that trigger a downstream cascade which results in the proposed inflammation and neurodegeneration observed in affected brains [3]. Although we have proposed an alternative hypothesis for the formation of Aβ-plaques and tau tangles in AD affected brain [4–6], both hypothesis conclude that abnormal protein accumulations may be due to a failing waste removal system in the aging brain. *Why have we been unable to identify the underlying cellular mechanisms for neurodegenerative diseases despite massive international research undertakings, advanced methodologies, and a vast body of AD-related publications?*

What is notably missing throughout this vast body of literature regarding waste removal from neurons are detailed structural studies, the basis of which inform a functional hypothesis. We have a solid understanding regarding structure and function of canal systems throughout the human body that fulfill the vitally important function of waste clearance. This includes the digestive-, urinary-, cardiovascular-, pulmonary-, and hepatic portal systems. What all these systems have in common is (*i*) the strict containment of waste products within a canal system from which the waste is sequestered and effectively expelled from the body, and (*ii*) waste-induced obstructions of these canal systems that may lead to life threatening conditions if untreated.

Currently, the prevalent ‘glymphatic hypothesis’ postulates that cellular waste is freely released from neurons into interstitial spaces throughout the brain and removed via the formation of an AQP4-mediated convective bulk flow of cerebrospinal fluid that flushes this debris from the brain parenchyma [7]. It has been suggested that the reduced flow of fluid in the aging brain leads to decreased efficacy of waste clearance and increased accumulation of misfolded protein in the brain parenchyma [7–10]. In an alternative postulation and compelled by the concept of an AQP4-mediated convective flow [7, 9] we have proposed the existence of a glial-canal system that internalizes cellular waste and transports debris toward the ventricular system [4–6]. Guided by our discovery of this canal system in giant neurons of the American Wandering spider *Cupiennius salei* we have postulated a novel role for myelin in waste removal [6]. According to the glial-canal hypothesis waste is strictly contained within this canal system and internalized via myelin-derived waste-canals that project into neuronal somata. We propose that cellular debris is drawn into waste-canals through an AQP4-mediated convective flow and channeled out of the brain parenchyma [6]. We postulate that obstructions of this canal system result in swelling of the AQP4-expressing glial cells which swell and appear spongiform. We propose that this swelling leads to rupture and uncontrolled depletion of associated neurons [4–6].

Here, we show the structural foundation of this proposed canal system in *C. salei* that has informed our functional hypothesis presented here. This arachnid model system is *exceptionally suitable* to study waste removal in healthy and degenerating neurons due to: (*i*) their susceptibility to neurodegenerative diseases that are manifested in behavioral abnormalities, (*ii*) the comparatively gigantic diameters of spider neurons (>100 µm) that allow us to unambiguously visualize the cellular structure and associated changes in neurodegeneration, (*iii*) clearly visible cellular waste that is up to 2 µm in diameter, (*iv*) easy accessibility, and (*v*) excellent tissue preservation. Moreover, sequencing of the spider genome has allowed us to show compelling homologies of key proteins that are implicated in neurodegeneration [11] between invertebrate and vertebrate species.

## MATERIALS

### Brain Tissue

#### Spider Brain

The data shown in this study were obtained from three healthy 12 to 14-month-old female spiders that showed no behavioral signs of neurodegeneration. The degenerating brain tissue was obtained from 18-, 21- and 36-month-old female spiders that showed early (18-month-old), progressing (21-month-old) and advanced (36-month-old) behavioral signs of neurodegeneration (see results).

#### Human Hippocampus

The representative human brain tissue shown here was obtained from a 79-year-old male decedent undergoing autopsy examination at the University of Vermont Medical Center. In compliance with Vermont State law the tissue was obtained with full consent for biomedical research, diagnosis, and teaching purposes from next of kin. All procedures were conducted in accordance with state laws. The hippocampal tissue sample was fixed in 4% paraformaldehyde with 2.5% glutaraldehyde. Dissection and diagnostic examination were carried out by Dr. John DeWitt (University of Vermont). Neuropathology showed significant accumulation of amyloid plaque and phosphorylated tau tangles in the hippocampus. The ABC score of A3B2C1 was consistent with intermediate burden Alzheimer disease neuropathologic change.

#### Rodent Brain

To prevent unnecessary animal sacrifices, we have utilized control rat tissue originating from healthy, 3-month-old male Sprague Dawley animals that were prepared for previous electron microscopic studies on rat brain. Animal use permissions and protocols strictly followed federal and state regulations.

Three mouse brains were obtained from surplus tissue from 16-week-old Cdh5-GCaMP8 wild type females at the University of Vermont. Although the brain tissue was surplus, we obtained IACUC approval prior to processing the brains.

### Tissue Preparation for Ultrastructural Analysis

Brain samples that were processed for electron microscopic investigations were fixed in a mixture of 4% paraformaldehyde (EMS 15710) and 2.5% glutaraldehyde (EMS 16019) in phosphate buffered saline pH 7.4, 0.1 M (PBS) overnight at 6°C. Dehydration and embedding in Araldite was performed as described previously [6]

### Semi- and Ultrathin Sectioning

Araldite-embedded brain specimens were cut using a Leica Ultracut E with an 8 mm Diatome histo-knife. To correlate structure at the light and electron microscopic levels section thickness was altered between 1-µm semithin and 60-nm serial ultrathin sections. Semithin sections were transferred onto glass slides, dried on a hot plate at 55 °C, stained with 2% aqueous toluidine blue (Sigma 6586-04-5) for ∼2 min at 55 °C, and rinsed thoroughly with distilled water.

### Toluidine-blue Stain of Vibratome Sections

We utilized 4% paraformaldehyde-fixed brain tissue to cut 70 µm vibratome sections with a Leica VT 1000S vibratome. After four 5 min wash cycles in cold PBS the sections were stained with a 2% aqueous toluidine blue solution. To avoid overstaining, we used a stereo dissection microscope. The 2% toluidine blue solution was slowly dripped into a petri dish containing the brain sections in PBS until the neurons appeared appropriately stained. The sections were embedded in Mowiol and examined promptly as the embedding medium will slowly de-stain the cells. The sections were examined using an Olympus microscope with differential phase contrast and digital image capturing capabilities.

### Immunohistochemistry

To ensure optimal tissue preservation the brain sections used for immunolabeling were kept on ice when possible. After fixation in 4% paraformaldehyde in PBS overnight at 6°C. The sections were washed in PBS prior to embedding in 4% Agarose (Sigma A9539). A Leica VT 1000S vibratome was used to cut 70-µm sections. The sections were transferred into 24-well dishes. After washing in PBS (4×5 min) we utilized 0.25% Bovine Serum Albumin (Sigma A4503) and 5% Normal Goat Serum (Sigma G9023) in 1% Triton-X/PBS to block unspecific binding sites (20 min). Incubation with the primary goat anti-rabbit AQP4 antiserum (BiCell #20104) and mouse anti-Myelin antibody (6-4H2 DSHB, IOWA) were conducted at dilutions of 1:100 in PBS containing 10% blocking medium overnight at 6°C. Specificity of AQP4-antibody binding in spider tissue has been described previously [6]. The sections were washed in PBS (5x 5 min) and immersed in blocking medium (20 min) prior to incubation with the secondary fluorochrome-coupled antibodies (Cy3 goat anti-rabbit, Jackson ImmunoResearch Laboratories 111-165-003, FITC goat anti-mouse Jackson ImmunoResearch Laboratories 115-096-072). The antibodies were used at a dilution of 1:600 in PBS containing 10% blocking medium overnight at 6 °C. To remove unbound antibodies the sections were washed in PBS, and incubated with Hoechst Blue nuclear stain (Sigma H 6024; 1:3000 in PBS) for 20 min. After washing in PBS for five additional wash cycles the sections were mounted on glass slides and embedded in Mowiol (Sigma# 81381). To avoid bleaching of the secondary antibodies all steps were conducted under minimum light conditions. The sections were analyzed using a confocal Zeiss AxioImager MZ with Apotome.

For control purposes and to establish the background fluorescence level of brain sections control preparations were processed under the omission of the primary antibody incubation. All samples examined were void of the fluorescent signals observed in sections treated with the primary antibodies.

### Luxol H&E Staining of Human Autopsy Tissue

Histological stains of human autopsy tissue were performed in the histology lab at the University of Vermont Health (formerly University of Vermont Medical Center) as described previously [6]. In brief: Brains were fixed in 10% neutral buffered formalin for at least 1 week at room temperature. Hippocampal tissue was embedded in paraffin, cut at 10 µm thickness and mounted on glass slides. Staining for myelin was condcted using 0.1% Luxol fast blue overnight prior to staining with Hematoxylin Eosin. Sections were permanently mounted using Permaslip Mounting Medium and Liquid Coverslip (Alban Scientific, Inc.).

### Tissue Sampling and DNA Extraction

We built a draft reference genome for *C. salei* and used it to identify and run a cross-species comparison of genes implicated in neurodegeneration. To build the genome, we sampled tissue from a 14-month-old healthy male spider. The animal was deeply anesthetized using CO^2^. The prosoma was submerged in cold 0.1M PBS pH 7 and the brain was removed and swiftly transferred into an Eppendorf tube that was frozen in liquid nitrogen. The tissue sample was transported in liquid nitrogen to the Vermont Integrated Genomic Resource at the University of Vermont (VIGR; RRID:SCR_021775) for sequencing and stored in a −80°C freezer. Tissue DNA was extracted using the E-Z 96 Tissue DNA Kit (Omega Bio-Tek, Norcross, GA). Long and short read sequences were generated from this tissue for genome assembly and annotation.

### Genome Sequencing

A Long-read sequencing library for direct whole genome DNA sequencing was generated using the Native Barcoding Kit (Oxford Nanopore Technologies (ONT), Oxford, UK) and was sequenced on the ONT PromethION 2 Solo device using R10.4.1 flow cell. We used Dorado version 0.7.2 to base call Pod5 files and then filter (min qscore 7; hac,5mCG_5hmCG), demultiplexed, and adapter trim the resulting sequences.

A short read whole genome DNA sequencing library was generated using the NEXTFLEX Rapid XP V2 DNA-Seq kit (Revvity, Waltham, MA). Library quality and concentration were determined using the High Sensitivity DNA Kit on Agilent Bioanalyzer 2100 and Qubit spectrofluorometer (ThermoFisher Scientific, Waltham, MA). The resulting library was paired-end (2x 150 bp reads) sequenced using the Singular Genomics G4 system (Singular Genomics, San Diego, CA).

### Genome Assembly and Annotation

Prior to genome assembly, the mitochondrial genome was assembled using GetOrganelle from Fastp quality trimmed and filtered short read sequence. The mitochondrial genome was standardized and annotated using MTGrasp. Before genome assembly, all long and short reads that mapped to the mitochondrial genome (using Dorado for long reads; minimap2 for short reads) were removed from the raw sequencing data using Samtools. We assembled the nuclear genome using the hybrid genome assembly nextflow workflow Pegasus. The resulting genome was annotated using the publicly available NCBI Eukaryotic Genome Annotation Pipeline. All steps of the DNA sequence processing and analysis were performed on the High-Performance Compute Cluster at the University of Vermont Advanced Computing Center (VACC; RRID:SCR_017762).

### Cross-species Comparison of Genes Implicated in Neurodegeneration

To investigate genes that are implicated in neurodegeneration, we surveyed the genome for the presence of genes related to amyloid-beta, aquaporins, the beta and gamma-secretase complex, microtubules, and myelin (Table 1). In cases where a gene of interest was not present in the genome annotation, we ran blast+ [12] searches against the draft genome using canonical human protein for that gene found in UniProtKB [13], which sources orthologous sequences from the Alliance of Genome [14] Resources, a collaborative effort that integrates comprehensive genetic and genomic data from multiple model organisms and humans. If a protein coding region identified in the draft genome annotation had multiple copies but one was complete, and the other was partial, we chose to use only the complete version in the analysis. We compiled fasta files of the protein products of those genes found in *C. salei, Homo sapiens, Mus musculus, Danio rerio, Drosophila melanogaster, and Caenorhabditis elegans.* The proteins for organisms (except *C. salei)* were downloaded from UniProt. We analyzed protein rather than nucleotide sequences to maximize the evolutionary signal across divergent taxa. Proteins were aligned using MAFFT [15] (alignment strategy set to auto), trimmed for evolutionarily relevant sites using ClipKIT [16] (default parameters), and maximum likelihood (ML) phylogenetic trees were built from the trimmed alignment using ModelFinder [17] in IQ-TREE 2 [18] with 1000 bootstrap replicates. We used R-Statistical Software (https://www.R-project.org/) to visualize multiple sequence alignments (R packages ggmsa [19] and ggplot2) and Phylogenetic trees (R packages ggtree and treeio [20]).

**Table 1.**
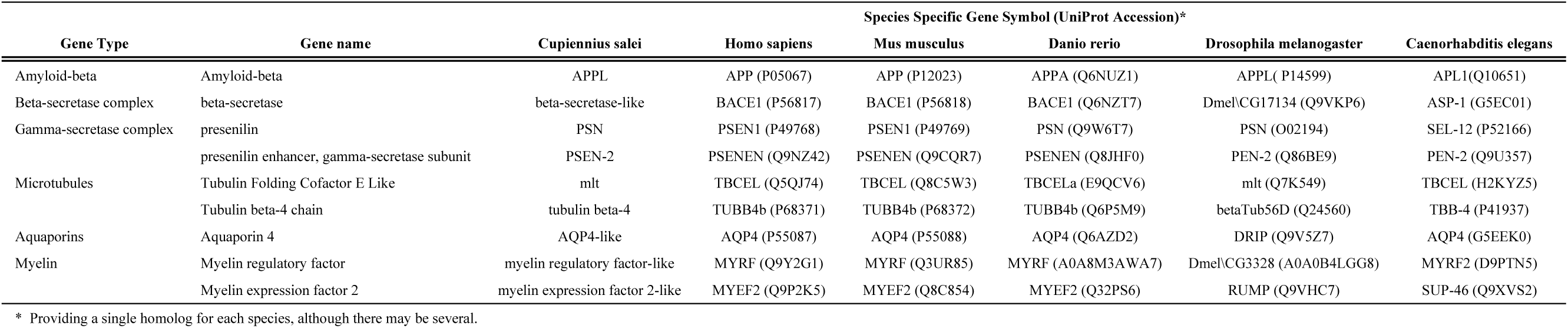
Homologous Alzheimer disease-associated genes found in C. salei, Human, and other model organisms.

### Image Processing

Confocal images were exported from the ZEN Blue program using the ‘Image export’ function. Figures were created using Adobe Photoshop.

### Glial Canal Measurements

The measurements of (*i*) waste-canals and (*ii*) glial sheaths were made using the measuring tool in ZEN Blue. Original images of Toluidine blue stained 1-µm semithin sections were captured using the x100 oil Objective. To ensure accuracy of the measurements, we worked with the original CZI-images that include metadata for the individual images.

To evaluate the size of (*i*) glial aqua-channels, (*ii*) neuronal aqua-channels, and (*iii*) aqua-canals in both healthy and degenerating spider tissue, blinded tissue sections of spider leg ganglia were evaluated to avoid bias. Captured EM images were measured in QuPath v0.6.0 [21] using appropriate scaling. To ensure correct identification, cellular features were manually selected and measured as polygons to determine area and the maximum diameter (i.e., major axis). While these data are entirely consistent with our prior work [6], it should be noted that we: (*i*) used a new collection of EM images for this study to confirm the robustness of our previous results, and (*ii*) switched our methodology from employing Fiji/ImageJ [22] to QuPath due to its project management feature.

### Statistical Analysis

Because all measured data were determined to be best fit by a log-normal distribution, multiple unequal variance, unpaired, two-tailed log-normal Student’s *t*-tests (with Holm-Šídák α-level adjustment for multiple comparisons) were conducted to perform comparisons of the cellular features across degenerating and healthy tissues. All statistical parameters and results are reported in the relevant figure legend. To the best of our knowledge, all assumptions of these tests were met. Statistics were calculated, and graphs were created using Prism 10.6.1 (GraphPad Software).

## RESULTS

### Behavioral Cues of Progressive Neurodegeneration in *C. salei*

Healthy adult (∼10-18 months) tropical wandering spiders rest on vertical surfaces head down. In this resting position both anterior prosoma and posterior opisthosoma remain in a straight line (Figure 1a). Histological investigation of healthy brains shows that giant motor neurons in the leg- and opisthosomal ganglia of these animals appear healthy and intact (Figure 1b). With increasing age (19 to 36-month-old), affected animals lose the ability to maintain their vertical resting position and their opisthosoma tilts forward with gravitational forces (Figure 1c). Toluidine blue-stained semithin sections of neurons in degenerating animals showed progressive depletion of the neuronal cytoplasm and spongiform swelling of surrounding glial cells (Figure 1d). To investigate cellular and ultrastructural changes in healthy compared to degenerating spiders, we have compared the neuroanatomy in healthy young adults with animals that showed (*i*) degeneration onset, and (*ii*) advanced degeneration.

**Figure 1.**
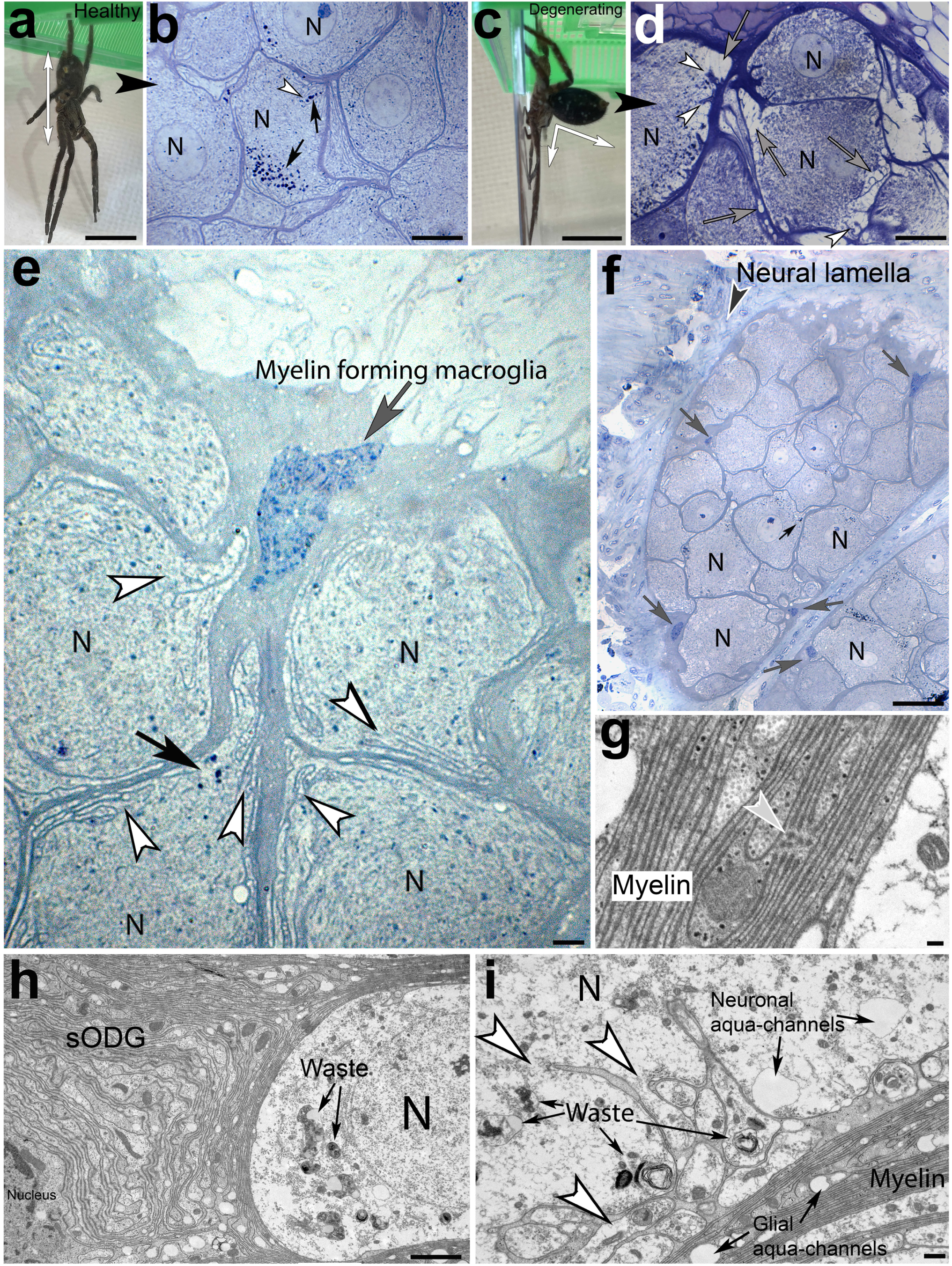
Neuroanatomy of healthy and degenerating giant neurons in the ventral leg ganglia of *Cupiennius salei*. **(a)** Healthy 24-month-old female in a typical vertical resting position shows straight alignment (*double headed arrow*) of the downward-facing prosoma and the upward-facing opisthosoma. **(b)** Toluidine blue-stained 1-μm semithin section through healthy neurons shows the typical accumulation of dark-blue stained lipofuscin particles (*black arrows*) within or around myelin-derived waste canals (*white arrowheads*) that partially detach from the glial sheath around the neuronal somata. The cytoplasm has a homogenous appearance, which is typical for healthy neurons. **(c)** Adult 24-month-old female with behavioral signs of advanced neurodegeneration. The upward-facing opisthosoma has a severe forward tilt in relation to the downward-facing prosoma (*white double arrow*). **(d)** Semithin section through neurons of degenerating brain tissue show hypertrophic swelling of the myelin sheath around neurons and waste canals that project perpendicular into the neuronal cytoplasm (*white arrowheads*). Spongiform abnormalities can be seen around the myelin-derived waste canals, and within the glial sheath (*grey arrows*). **(e-f)** Semithin sections through healthy brain tissue show an interconnected network of heavily branching macroglia (*grey arrows*) whose somata are located along the outer periphery of the ganglia adjacent to the neural lamella. The partial detachment of finger-like myelin canals that project into the neuronal somata (*white arrowheads*) is most prominent in neurons that border on the somata of macroglia. **(g-i)** Electron micrographs show the myelinated nature of macroglia and intraneuronal waste canals that engulf cellular waste. The myelin sheath contains ‘microtubule-associated breaking points’ where individual myelin membranes appear severed. The membrane endings are closely associated with a cluster of five microtubules (*grey arrowhead in g*). Numerous electron-lucent profiles of ‘glial aqua-channels’ are contained in the myelin sheath, whereas intraneuronal waste-canals (*white arrowheads in i*) are near neuronal aqua-channels with electron-lucent lumina (*panel i*). (*N*): Neuronal somata. Scale bars: a: 1.5 cm; b: 20 µm; c: 1.5 cm; d: 20 µm; e: 5 µm; f: 50 µm; g: 100 nm; h: 2 µm; i: 500 nm.

### Myelin-forming macroglia give rise to waste-internalizing canals in healthy spider brain

Neurons in healthy spider brain are ensheathed by a prominent myelin sheath (Figure 1).

Histological investigation of toluidine blue-stained semithin sections and electron microscopic ultrathin sections show that this sheath originates from a syncytium of macroglia referred to as spider oligodendroglia (sODG; Figure 1). The somata of these macroglia reside in the ‘neural lamella’ that surrounds each ganglion and borders on the lymphatic system (Figures 1, 2). Long myelin processes project into the brain ganglia and give rise to waste-canals that project into neurons and internalize cellular waste (Figures 1-3). In the following we will refer to these structures as ‘waste-canals.’ The myelin sheath around neuronal somata that are adjacent to sODG can be >10 µm thick (Figure 1e, f, h). In contrast, the myelin sheath around medially located neurons ranges between ∼0.3 −4 µm in thickness at the light microscopic level and can be as narrow as 200 nm consisting only of ∼10 myelin membranes at the ultrastructural level (Figure 2). Interestingly, the membranes within the myelin sheath are not continuous but contain ‘microtubule-associated breaking points’ (MABs). In these regions the myelin membranes appear cleaved whereby the cleaved membranes appear ‘anchored’ to clusters of ∼1-12 microtubules (Figure 1g). Around the neuronal cortex, where glial cells and neurons border on each other, numerous glial lobes partially detach and project into the neuronal somata where they form lipofuscin-internalizing waste-canals (Figures 2-4).

**Figure 2:**
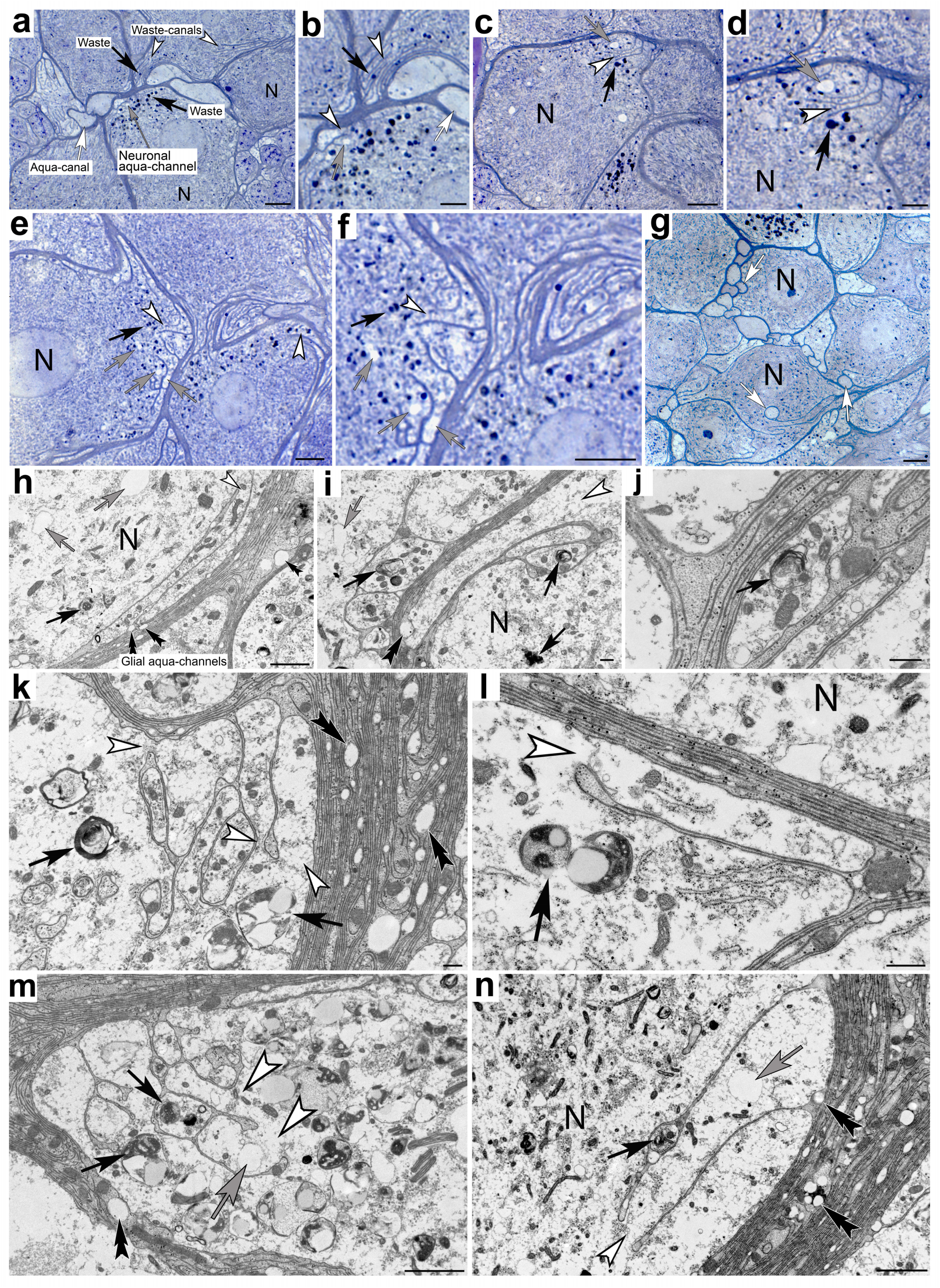
Histological and ultrastructural depiction of (*i*) waste-canals, (*ii*) aqua-canals, (*iii*) neuronal aqua-channels and glial-aqua-channels in healthy leg ganglia of *C. salei*. **(a-g)** Toluidine blue-stained semithin sections Neuronal somata (*N*) are surrounded by macroglia processes from which finger-like glial lobes partially detach and project into the neuronal somata *(white arrowheads*). Large diameter electron lucent aqua-canals can be observed adjacent to (*a, b*), or within the neuronal cytoplasm (*white arrows in g*). Lipofuscin granules (*black arrows*) accumulate near waste-canals (*white arrowheads*), which are frequently observed to detach in an annular fashion from the myelin sheath that surrounds aqua-canals (*b*). Neuronal aqua-channels (*grey arrows*) are observed near waste-canals which explains the slightly electron-lucent appearance in these areas. **(h-n)** Ultrastructural depiction of waste-canals demonstrates the waste-internalizing nature of the myelin-derived canals that is consistent with light microscopic observations. Neuronal aqua-channels can be seen around and within the lumina of individual waste-canals (*grey arrow in n*). Glial aqua-channels (*black double arrowheads*) are abundant in the myelin sheath. *White arrowheads*: waste-canals; *black arrows*: cellular waste; *grey arrows*: neuronal aqua-channels; *black arrowheads*: glial aqua-channels. Scale bars: a: 10 µm; b: 5 µm; c: 10 µm; d: 5 µm; e-g: 10 µm; h: 2 µm ;i-l: 500 nm.

**Figure 3.**
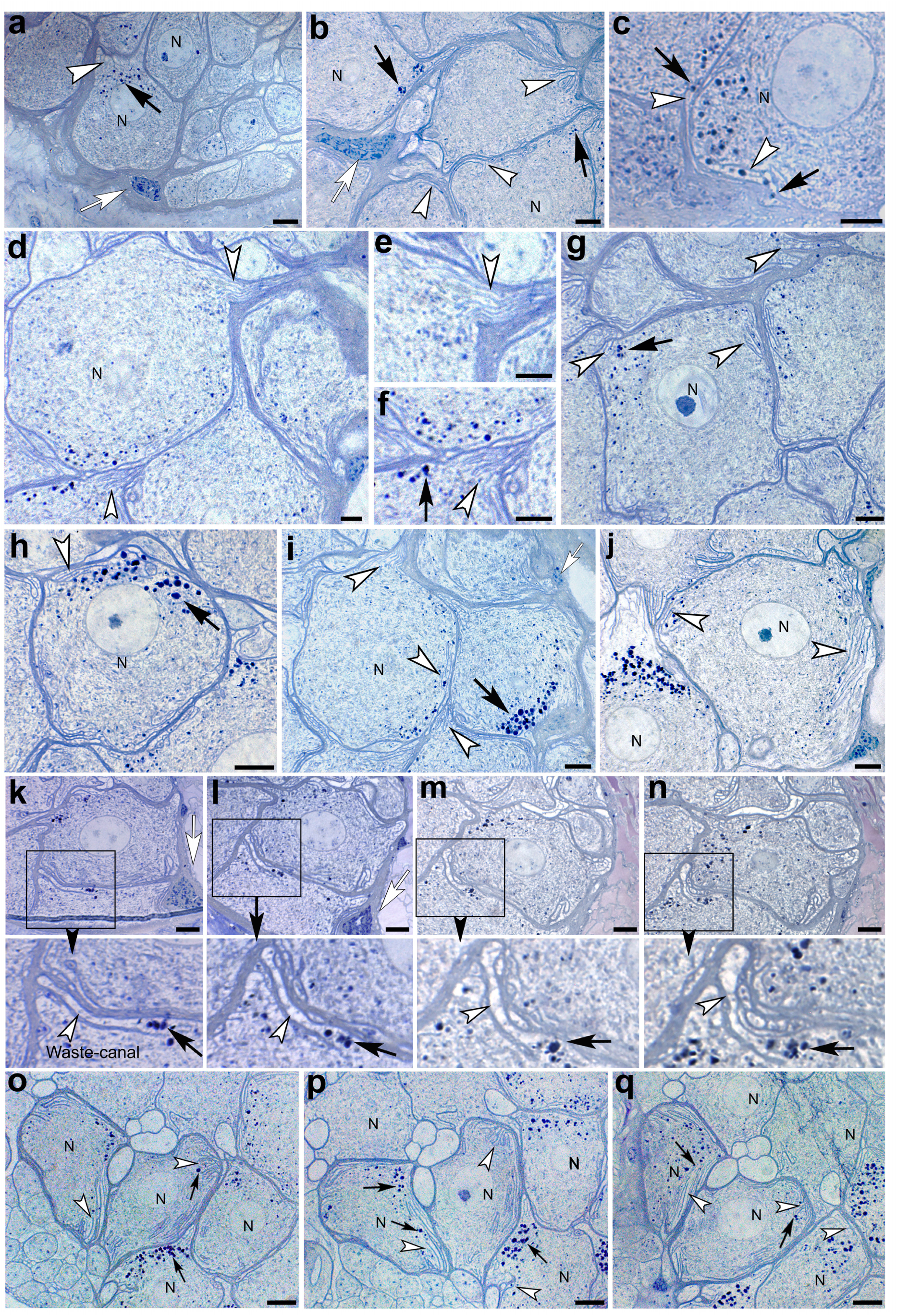
Toluidine blue-stained single and serial sections of healthy giant spider neurons in the leg ganglia of *C. salei*. **(a-j)** Cross-sections through neuronal somata demonstrate the abundance of glia-derived waste-canals that partially detach from the surrounding glial processes numerous lipofuscin particles can be seen accumulating around and within the waste-canals. **(k-d)** Two sets of serial sections through individual neurons (*k-n with higher zoom insets below and o-q*) demonstrate the consistency by which waste-canals are formed and internalize cellular waste. *White arrowheads*: waste-canals; *black arrows*: cellular waste; *grey arrows*: neuronal aqua-channels. Scale bars: 10 µm

**Figure 4.**
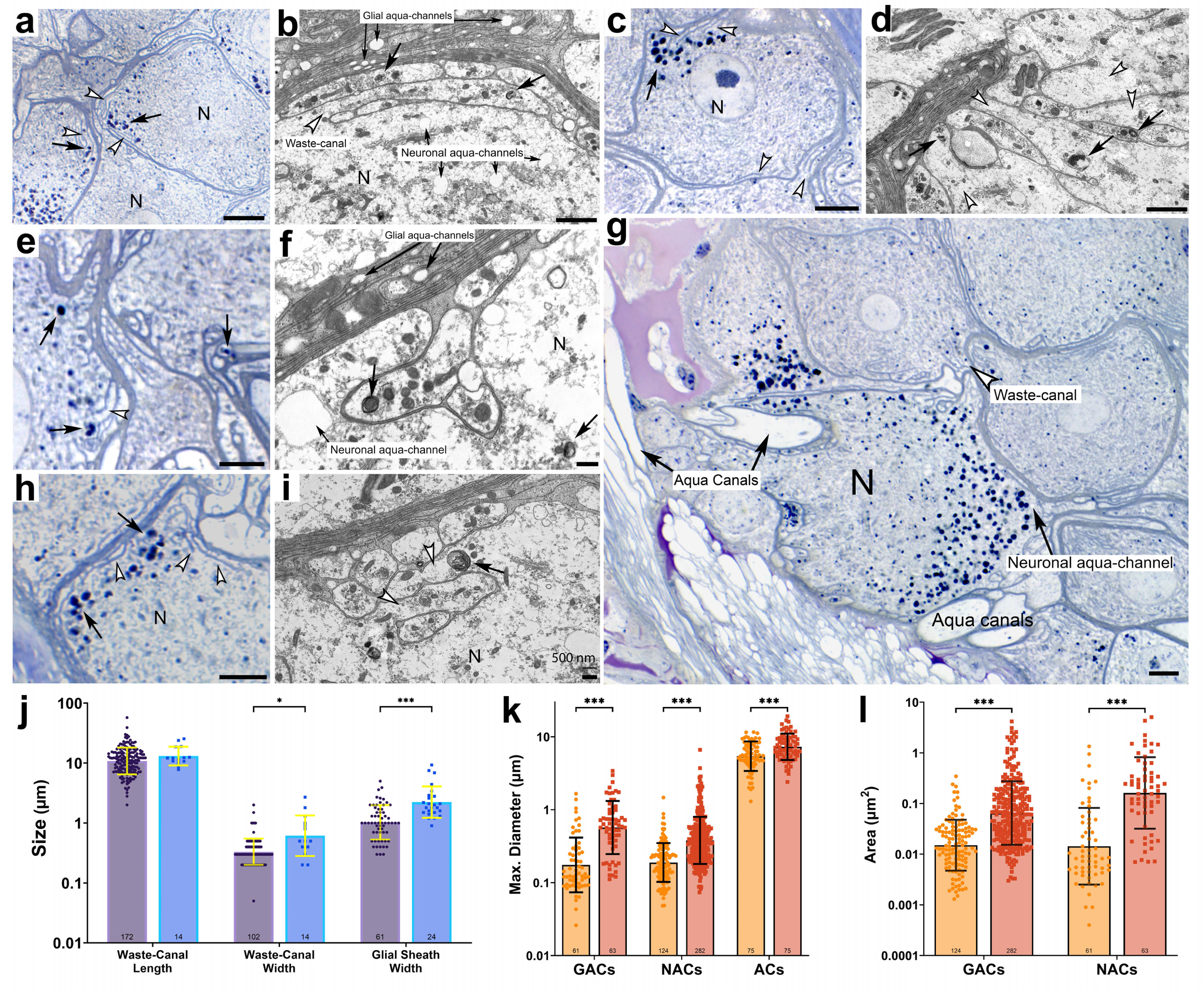
Light- and electron microscopic depiction of waste canals in giant spider neurons. **(a, c, e, g,h)** Toluidine blue-stained semithin sections show the abundance of lipofuscin (*black arrows*) within and around myelin-derived waste canals (*white arrowheads*). Aqua canals with clear lumina project from the lymphatic system into adjacent neurons (*I*). **(b, d, f, i)** Electron micrographs show the ultrastructure and diversity of waste-internalizing myelin-derived waste-canals and their association with neuronal- and glial aqua-canals. **(j)** Average length of the waste-canals is not significantly different in degenerating (13.1 μm GM) compared with healthy tissue (10.8 μm GM; *t*(17.6) = 1.82, *p* = 0.085, unpaired *t*-test). Average width of the waste-canals is significantly larger in degenerating (0.614 μm GM) compared to healthy tissue (0.332 μm GM; *t*(14.5) = 2.88, *p* = 0.023, unpaired *t*-test). Average width of the glial sheath is significantly larger in degenerating (2.23 μm GM) compared to healthy tissue (1.03 μm GM; *t*(46.1) = 5.2, *p* < 0.001, unpaired *t*-test). **(k)** Average maximum diameter of the GACs is significantly larger in degenerating (0.570 μm GM) compared to healthy tissue (0.175 μm GM; *t*(121.5) = 7.720, *p* < 0.001, unpaired *t*-test). Average maximum diameter of the NACs is significantly larger in degenerating (0.379 μm GM) compared to healthy tissue (0.189 μm GM; *t*(282.2) = 9.864, *p* < 0.001, unpaired *t*-test). Average maximum diameter of the ACs is significantly larger in degenerating (7.287 μm GM) compared to healthy tissue (5.422 μm GM; *t*(146.2) = 4.101, *p* < 0.001, unpaired *t*-test). **(l)** The average area of the GACs is significantly larger in degenerating (0.0648 μm^2^ GM) compared to healthy tissue (0.015 μm^2^ GM; *t*(289.6) = 10.84, *p* < 0.001, unpaired *t*-test). The average area of the NACs is significantly larger in degenerating (0.162 μm^2^ GM) compared to healthy tissue (0.014 μm^2^ GM; *t*(120.8) = 8.005, *p* < 0.001, unpaired *t*-test). Graph symbols: healthy(●), degenerating (▪). Significance: *: p < 0.05; **: p < 0.01; ***: p < 0.001.

Both semi- and ultrathin sections revealed three conspicuous structures that are associated with waste-canals. For clarity, we have labeled these structures in Figure 2. Firstly, large diameter canal-like cell processes with diameters between 1.3 and 11.6 µm in the following referred to as ‘aqua-canals’ (ACs; Figure 2a). The lumina of ACs are void of cytoplasmic texture and appear clear in both cross and longitudinal section planes. Interestingly, the myelin sheath around ACs frequently detaches and forms waste-canals both along the outer and inner surfaces. Due to the circular structure of ACs the detaching waste-canals have an annular appearance (Figure 2b). The second structures that are observed in association with glial canals are smaller ‘neuronal aqua-channels’ (NACs; Figure 2). Like ACs, the lumina of NACs appear void of cytoplasmic structure at both the light- and electron microscopic levels. Surrounded by a cell membrane, NACs are located within the neuronal cytoplasm near waste-canals (Figure 2). Ultrastructural investigations show that swelling NACs are often located within waste-canals (Figure 2m, n). Due to the presence of NACs near waste-canals the neuronal cytoplasm in and around waste-canals appears lighter compared to cytoplasm that is distant from waste canals (Figure 2a-f). In healthy brain tissue, the diameters of NACs vary between 0.048 and 1.468 µm.

The third conspicuous structures are translucent channel-like structures that are located within the myelin sheath (Figure 2h-n). We will refer to these structures as ‘glial aqua-channels’ (GACs). In healthy brain tissue, the diameters of GACs are ∼250 nm but can range between 0.026 and 1.659 µm.

### Structural characterization of lipofuscin and waste-canals in healthy spider brain

The diameters of lipofuscin that accumulates in areas where waste-canals form vary between ∼ 0.1 µm to >2 µm. At the ultrastructural level, this cellular waste consists of a variety of lamellar accumulations and electron-dense particles (Figures 1, 2). A subpopulation of lipofuscin particles exhibits electron-lucent areas (Figure 1h, 2l). Individual myelin membranes partially detach from the main glial sheath and give rise to one or more finger-like canals that project into the neuronal cytoplasm and internalize both lipofuscin and organelles (Figures 1, 2, 4). These canals are parallel to either the neuronal perimeter or adjacent detaching glial lobes, which leads to the formation of one or more narrow canals (Figure 3j). Size and shape of individual waste-canals varied widely. Whereas some consisted of a single myelin-derived cell membrane (Figure 2l, n), others ranged between ∼2-57 µm in length and ∼0.05-2 µm in width (Figures 2, 3, 4j). Larger waste-canals often appeared to originate from one common glial lobe whose apical end separates into 2-11 finger-like canals (Figures 1-4). Internalized lipofuscin can also be observed within the myelin sheath around neuronal somata (Figure 2j). Serial section analysis demonstrates the prominent lipofuscin accumulations particles near waste canals (Figures 3k-q, S1).

### Immunolabeling for AQP4 reveals immunoreactivity in areas consistent with the location of ACs

To test the hypothesis that the accumulation of lipofuscin near waste-canals may be due to an AQP4-mediated convective flow we have conducted immunolabeling for AQP4. Observed punctate immunolabeling within and around neuronal somata is consistent with size and location of the above-described canal structures. Interestingly, the immunofluorescence appears both circular and longitudinally consistent with putative canal structures, whereas the sODG appears immuno-negative (Figures S1, 5).

### Spongiform pathologies in degenerating spider brain are caused by hypertrophic swelling of GACs and NACs

To conduct comparative investigations of the neuroanatomy in degenerating spider neurons we have examined Toluidine blue-stained semithin sections through spider brains of 18- and 36-month-old animals. The 18-month-old animals (*n=3*) showed mild signs of neurodegeneration. The 36-month-old animals (*n=3*) showed severe behavioral signs of neurodegeneration.

Histopathological signs of early neurodegeneration were hypertrophic swelling of the glial sheath and intraneuronal glial projections that contribute to the formation of waste canals (Figure 5a-c). These structural pathologies coincided with comparatively mild spongiform abnormalities and unraveling of the glial sheath around neuronal somata. The latter caused neurons to appear ‘zig-zag’ shaped (Figure 5).

**Figure 5.**
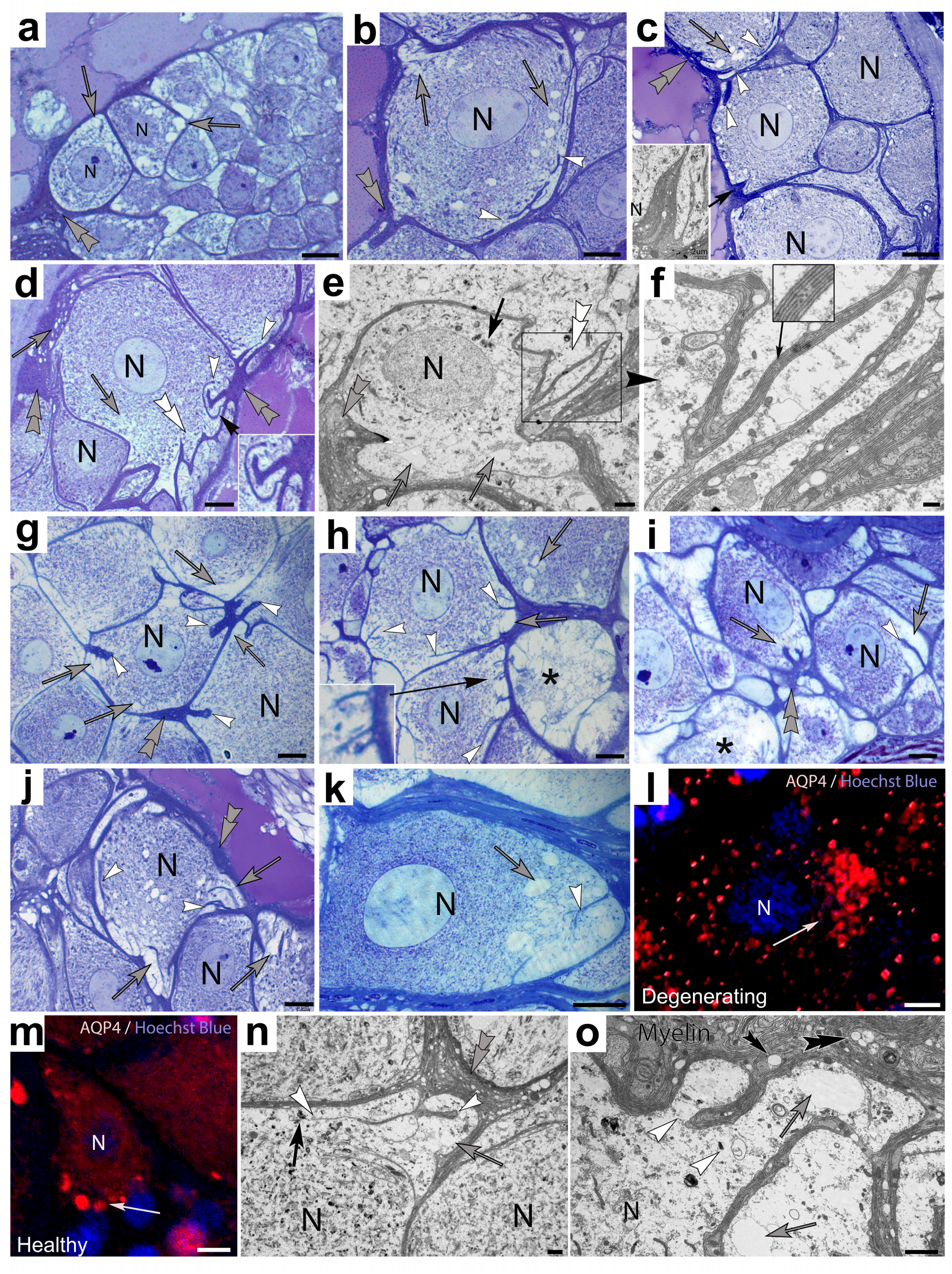
Histopathological changes in degenerating leg ganglia of *Cupiennius salei*. **(a-g)** Light- (*a,b,c,d,g*) and electron microscopic (*e, f*) depiction of neurons with early signs of neurodegeneration reveals (*i*) hypertrophic swelling of the glial sheath (grey double arrowheads), (*ii*) beginning spongiform abnormalities within the glial sheath and around the periphery of neuronal somata and waste-canals (white arrowheads), and (*iii*) unraveling of the glial-sheath. **(h-k)** Toluidine blue-stained neurons in advanced neurodegenerative stages demonstrate the progressing spongiform depletion of cytoplasmic content that progresses toward the medial areas of neurons, ultimately depleting the cells of their cytoplasmic content (*asterisks*). Please note the advanced spongiform pathologies in the vicinity of waste-canals (*white arrowheads*) compared to cytoplasm that is void of the latter. **(l, m)** Immunolabeling for AQP4 in degenerating (*l*) compared to healthy neurons (*m*) shows increasing immunoreactivity within the neuronal somata in advancing degeneration (*white arrow in l*) consistent with the location of spongiform depletion in toluidine blue stained neurons (*grey arrow in k*). **(n, o)** Ultrastructural depiction of degenerating neurons demonstrate the hypertrophic swelling of neuronal- (*grey arrows*) and glial aqua-canals (*black double arrowheads*) consistent with the spongiform pathologies observed at the light microscopic level, myelinated glial canals disassociate from the microtubules and project into the neuronal somata creating an opening (*white arrowhead*) with waste (*black arrow*) entering canal. Scale bars: a-d: 10 µm; e: 2 µm; f: 500 nm; g-j: 10 µm; k: 20 µm; l: 5 µm; m: 10 µm; n-o: 2 µm.

Both maximum diameter and area of individual NACs were significantly larger in degenerating (0.570 μm and 0.1617 μm^2^ geometric means [GM], respectively) compared to healthy tissue (0.175 μm and 0.0143 μm^2^ GM; Figure 4) indicative of swelling NACs. Similarly, both maximum diameter and area of individual GACs were significantly larger in degenerating (0.379 μm and 0.0648 μm^2^ GM, respectively) compared to healthy tissue (0.189 μm and 0.0151 μm^2^ GM; Figure 4), indicative of swelling GACs. Finally, the maximum diameter of individual ACs were significantly larger in degenerating (7.29 μm GM, respectively) compared to healthy tissue (5.42 μm GM; Figure 4), consistent with swelling of ACs.

Due to the translucent lumina of these swelling canals affected tissue appears spongiform, which is particularly conspicuous around waste-canals where GACs swell to diameters of up to 12 µm (Figure 5k). Similar swelling was observed in GACs within the myelin sheath (Figure 5d, n), which explained the increasingly spongiform appearance of degenerating brain tissue (Figure 5). Swelling GACs can often be seen emanating from hypertrophic myelin lobes of waste canals (Figure 5g, j). As a result, the cytoplasm in the vicinity of waste-canals is replaced by swelling NACs and canal-forming glial lobes are no longer parallel to the glial sheath but appear perpendicular to the neuronal perimeter (Figure 5). Lipofuscin granules were largely absent around degenerating waste-canals. Immunolabeling of degenerating brain tissue demonstrates that increased AQP4-immunoreactivity within degenerating brain tissue is consistent with the light microscopic appearance of hypertrophic GACs throughout the neurons and in the vicinity of waste-canals (Figure 5k, l). In contrast, AQP4-immunoreactivity in healthy brain tissue appeared more restricted to cell profiles whose appearance was consistent with ACs (Figure 5m).

### Comparison of myelinated glial cells in spider, rodent, and human brain

Comparison of myelin-forming glial cells in leg ganglia of *C. salei* with the hippocampal formation of *Rattus norvegicus* and *homo sapiens* demonstrates intriguing parallels. In all three species heavily, myelinated cells reside in the periphery of the brain adjacent to the neural lamella (*C. salei;* Figure 6 a-c) or ventricles (*R. norvegicus,* Figure 6 d-g*; H sapiens,* Figure 6 o-x). In all three species the myelin-forming glial cells form long varicose projections into the brain parenchyma that make close contact with neurons (*C. salei*: Figures 1e, 6b, c; *R. norvegicus*: Figure 6e, i; *H. sapiens*: Figure 6p-x). Both light- and electron microscopic investigations suggest that in all three species myelin-derived processes transect into neuronal somata and internalize cellular waste known as lipofuscin in *C. salei*, (Figure 2) or electron-dense material in *R. norvegicus* (Figure 6 k-n), and *H. sapiens* (Figure 6q-x). Interestingly, investigation of rat hippocampal axons that were unambiguously identified as axonal processes based on their physical association to neuronal somata lacked myelination even well beyond the axon hillock (Figure 6h). However, myelinated, varicose cell profiles whose narrow (0.5-1 µm) diameters are consistent with the cell processes formed by ependymal tanycytes were seen to project parallel to the larger axon profiles and form varicose projections into the axonal processes (Figure 7h, i). In all three species AQP4-immunoreactive cell profiles can be observed in cell profiles and locations consistent with myelinated tanycyte processes (*C. salei*, Figures S1, 5; *R. norvegicus*, Figure 6 j; *H. sapiens*, Figure 6 o). Both light and ultrastructural investigation of myelinated cell processes in AD affected human brain show the emergence of myelin-derived receptacles that project into neuronal somata (Figure 6r-x). At the ultrastructural level, these receptacles contain electron-dense material consistent with the appearance of cellular waste in mammals.

**Figure 6.**
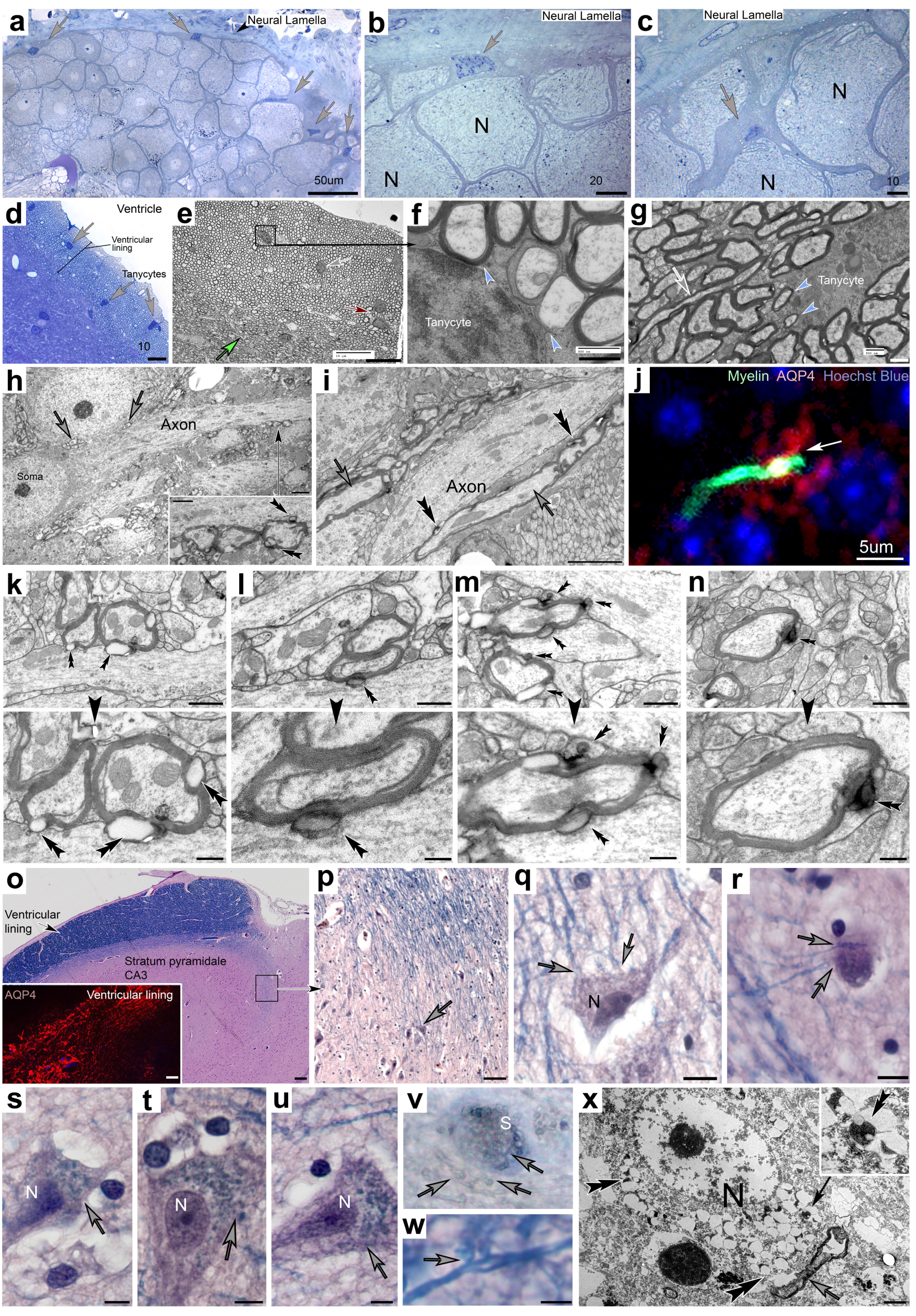
Comparison of myelinated glial cells in the brains of *Cupiennius salei*, *Rattus norvegicus* and *Homo sapiens*. **(a-c)** Leg ganglia in *C. salei* are surrounded by a neural lamella that borders the lymphatic system. The somata of myelin-forming macroglia are located adjacent to the neural lamella and send a vast network of myelinated processes into the underlying brain parenchyma surrounding neuronal somata with a myelin sheath. **(d-g)** The hippocampal alveus in *R. norvegicus* borders on the third ventricle and contains a syncytial network (*red arrow in e*) of ependymal tanycytes (*grey arrows*) that form a vast network of myelinated processes. Forming myelinated processes can be seen within tanycyte somata (*blue arrowheads*). The myelinated cell processes project into the underlying brain parenchyma and start to form numerous electron-lucent varicosities (*green arrow in e*). **(h, i)** Electron micrographs of rat hippocampus show myelinated varicosity-forming (*black double arrowheads*) cell profiles (*grey arrows*) project parallel to the axon of a hippocampal pyramidal cell. **(j)** Double labeling for myelin and AQP4 of mouse hippocampus reveals colocalization of both epitopes, whereby the AQP4-immunoreactive processes can often be seen to emanate from the myelin-immunoreactive cell process (*arrow in j*). **(k-n)** Varicose processes that protrude from myelinated cell profiles and project into adjacent cell processes appear either electron lucent (*black double arrowheads in k, inset below*) or contain varying amounts of electron-dense material consistent with cellular waste in mammalian brain (*black double arrowheads in l-n, insets below).* **(o-u)** Luxol blue/Hematoxylin Eosin-stained human hippocampus reveals a vast fiber network that is stained for the myelin marker Luxol blue in the ventricular lining (*o*). Fine cell processes project from the alveus into the underlying stratum pyramidale where the fibers project into cell somata (*grey arrows*) and form varicose receptacles (*grey arrows s-u*). **(v)** Toluidine blue 70-µm vibratome sections show the dense obstruction of a soma (*s*) with bulging varicosities that are consistent with varicosities observed on cell processes adjacent to the soma (*grey arrows in v*). **(w)** Luxol H&E stained varicose tanycyte process in Alzheimer affected human brain tissue gives rise to varicose receptacles (*grey arrow*) **(x)** Ultrastructural depiction of an Alzheimer affected neuron (*N*) and adjacent myelinated varicosity-forming cell process (*grey arrow*). The latter forms numerous varicosities (*black double arrowheads, Inset*) that project into the neuronal soma. The appearance of these varicosities is consistent with human lipofuscin (*inset*). Scale bars: a: 50 µm; b: 20 µm; c-e: 10 µm; f, g: 500 nm; h: 2 µm, Inset: 500 nm; i: 2 µm ; j: 5 µm; k-n, Insets: 500 nm, o: 200 µm, Inset: 20 µm; p: 50 µm, q, r: 10 µm; s-u: 5 µm, v: 10 µm; w: 2.5 µm, x: 2 µm.

**Figure 7.**
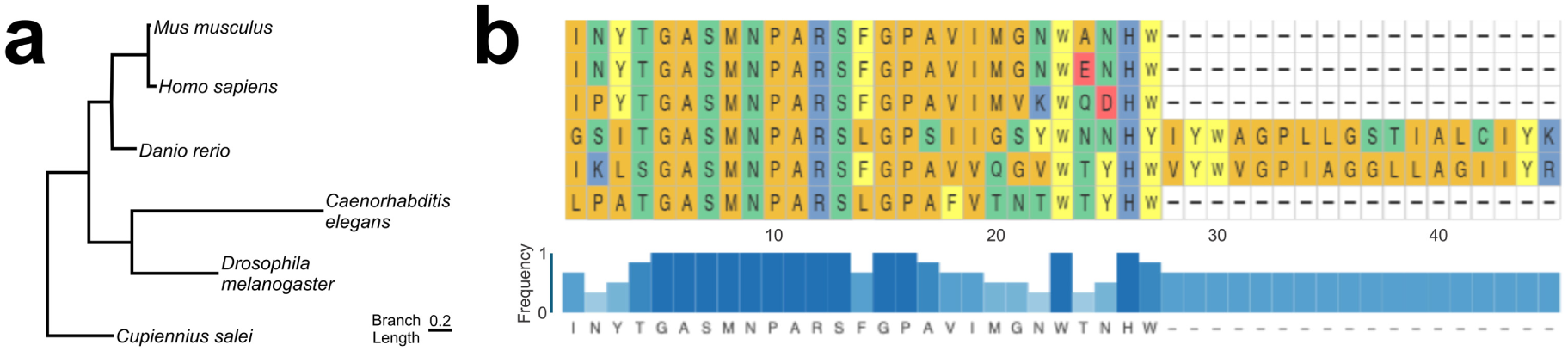
Comparison of AQP4 across model organisms. (**a)** A Maximum likelihood tree of the entire protein coding regions of AQP4 for *Cupiennius salei, Homo sapiens, Mus musculus, Danio rerio, Caenorhabditis elegans,* and *Drosophila melanogaster* generated from amino acids. **(b)** Amino acid alignment and consensus sequence for the conserved region of AQP4 that corresponds with Exon 4 in the canonical Homo sapiens protein (UniProt Accession P55087).

### Sequencing of the C. salei genome reveals homologues for the human AQP4 gene and other AD-related genes

The draft genome generated for *C. salei* is comprised of a mitochondrial (NCBI accessions pending) and nuclear genome (NCBI accessions pending). The draft mitochondrial genome is circular, 14,738 bp long, and has an average short read base coverage of 1659.4x and a k-mer coverage of 156x. It contains 13 protein-coding genes, 21 transfer RNAs, and 2 ribosomal RNAs. The nuclear genome is 2.51 Gbp in length with a GC content of 31.6 and consists of 12785 scaffolds and an N50 of 2.8 Mbp. The mean short read coverage of the genome is 8.2x and the mean long read coverage is 25x. The Pegasus workflow calculated the number of Universal Single-Copy Orthologs present in the genome build based on what is known from arachnid genomes, using BUSCO version 5.4.7 arachnida_odb10 database. The *C. salei* draft genome assembly contains 98.4% complete BUSCOs (n=2934 BUSCO groups; 2.7% of the expected complete BUSCOs are single copy, 5.7% of the expected complete BUSCOs are duplicated), 0.3% of the BUSCOs are fragmented, and 1.3% of the BUSCOs are missing. Annotation of the draft genome using EGAPx determined it contains 14,231 genes, and 18,894 mRNA transcripts. Our findings reveal that the *C. salei* genome is comprised of 0.93-6+ Gb.

### Cross-species Comparison of Homologous Genes

Our search for genes implicated with AD showed numerous homologous genes present in the draft *C. salei* genome (Table 1, Figures S2, 3) including AQP4, Aβ, β-secretase, presenilin, presenilin enhancer, gamma-secretase subunit, Tubulin Folding Cofactor E Like, Tubulin beta-4 chain, Myelin regulatory factor, Myelin expression factor 2. Although there is sequence divergence between species, there are conserved regions shared by all species (Figures S2, 3). For example, we present a phylogenetic tree and multiple sequence alignment of conserved sites for AQP4 (Figure 7).

The AQP4 complete multiple sequence alignment used for phylogenetic analysis had 325 characters, 305 distinct patterns, 79 parsimony-informative sites, 193 singleton sites, and 52 constant sites. The best-fit model of amino acid sequence substitution was Q.PFAM+G4 and chosen according to Bayesian Information Criterion. The single ML tree had a log-likelihood of - 3362.710 and the ML bootstrap consensus tree gave the same topology as well as confidence values for each clade (Figure 7a). We identified two conserved regions in *C. salei* AQP4: two discontinuously helical intramembrane regions (GHI**NPA**VT (*Homo sapiens* exon2) and GASM**NPA**RSFGPAV (*Homo sapiens* exon4; Figure 7b) with two conserved NPA motifs (asparagine-proline-alanine, NPA). These regions are conserved across a broad phylogenetic range as seen in the AQP4 multiple alignment with *C. salei*, *Homo sapiens*, *Mus musculus*, *Danio rerio*, *Drosophila melanogaster* and *Caenorhabditis elegans (Figure 7b)*.

### Proposed functional model of the myelin-derived waste-canal system

Based on the findings presented here, we propose the existence of a myelin-mediated glial-canal system that internalizes cellular waste in an AQP4-dependent manner that may be highly conserved in the mammalian brain. We postulate that myelinated macroglia provide cell membranes for the sustainable formation of waste-internalizing canals (Figure 8a). We postulate that these canals contain APQ4-expressing channels that contribute to the formation of a convective bulk flow that channels waste into the canal-system (Figure 8). We postulate that pathological swelling of the AQP4 expressing cell processes leads to spongiform hypertrophy of affected cell profiles and associated neurons that result in depletion and death of affected brain tissue (Figure 8b).

**Figure 8.**
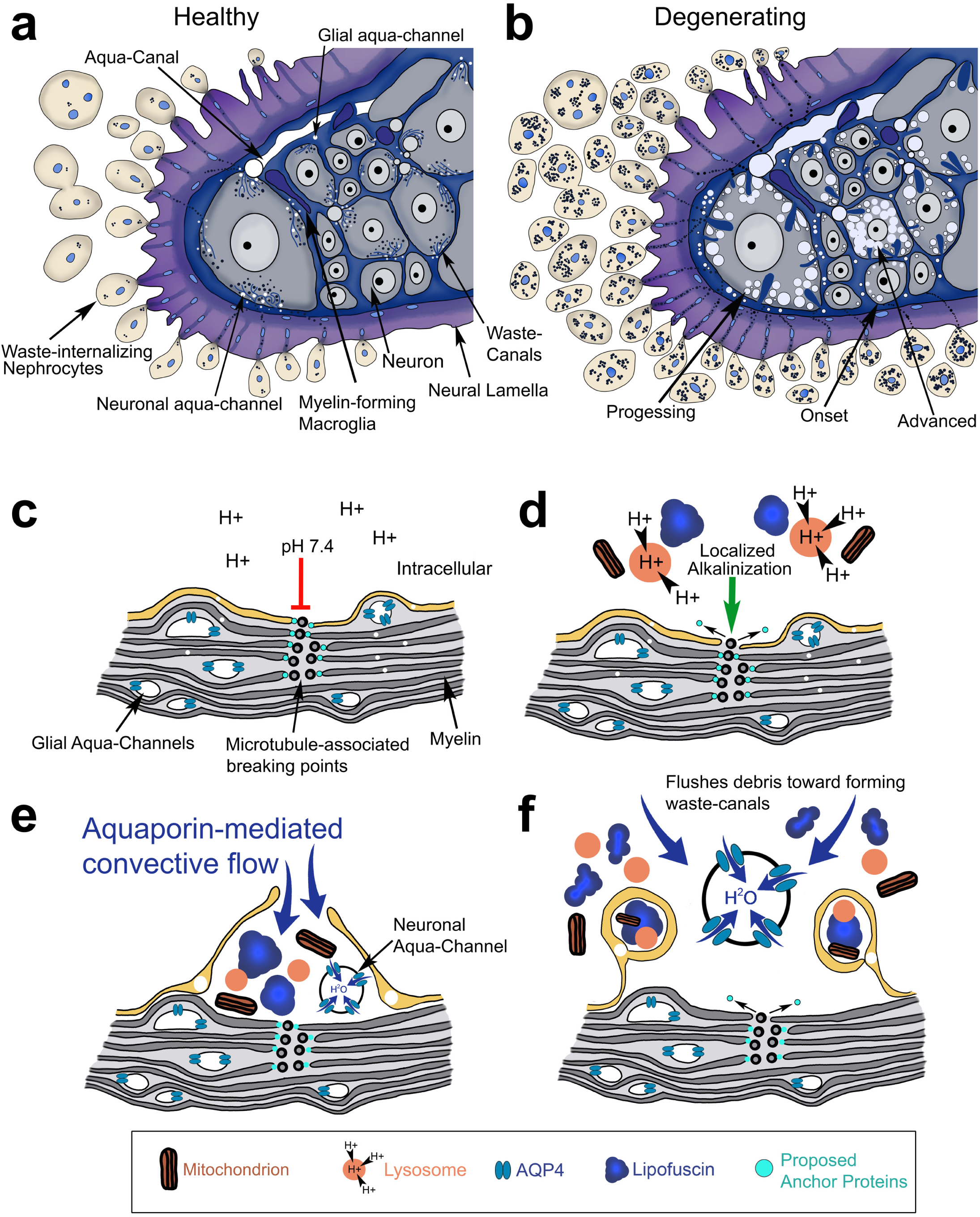
Schematic depictions of the proposed functional model of waste removal from spider neurons in health and degeneration. **(a)** Healthy brain tissue: Giant neurons in ventral leg ganglion are surrounded by myelin-forming macroglia that give rise to debris-internalizing waste-canals that project into the neuronal cytoplasm. Numerous neuronal aqua-channels are present in the vicinity of waste-canals. Glial aqua-channels are contained in the myelin sheath. Internalized waste is channeled through canals in the neural lamella and internalized by mesenchymal nephrocytes that fuse into multinucleated syncytia. **(b)** Degenerating brain tissue: Myelinated glial cells, aqua-canals, glial- and neuronal aqua-channels show hypertrophic swelling that leads to mild spongiform appearance around the neuronal periphery during degeneration *onset*. Neurons with *progressing* signs of degeneration show prominent spongiform swellings around waste-canals and the cellular cortex. In *advanced* degeneration the neuronal cytoplasm is largely replaced by spongiform abnormalities. Increased numbers of waste-containing macrophages emerge from the neural lamella. **(c-f)** Working hypothesis regarding the functional significance of microtubule-associated-breaking points (MABs) in the formation of myelin-derived waste-canals. (*c*) In the absence of cellular waste, the neutral cytoplasmic pH 7.4 inhibits (*red bar*) the dissociation of MABs, cleaved myelin membranes remain anchored to microtubules. (*d*) In the presence of waste lysosomal proton pumps cause localized alkalinization. The change in pH promotes (*green arrow*) detachment of myelin membranes from their microtubule anchors. (d) translocation of glial aqua-channel into neuronal cytoplasm activates aquaporin4 water channels and creates a convective cytoplasmic flow into the waste-canal. (f) Waste together with organelles required for catabolism are sequestered from the neuronal cytoplasm and channeled out of the brain via lymphatic system. Neuronal aqua-channel creates convective flow toward MABs explaining (*i*) the formation of numerous waste canals within a given area, and (*ii*) accumulation of waste near waste-canals. Figure not drawn to scale.

We postulate that the physiological formation of waste canals in healthy brain is governed through the controlled release of myelin membranes via MABs. This process may be governed through lysosome-induced localized alkalinization in the presence of cellular debris that may trigger the enzymatic release of anchor proteins from microtubules (Figure 8c-f).

## DISCUSSION

This is the first study that provides a detailed structural foundation of a recently proposed glial derived waste-canal system in giant motor neurons of *C. salei*. We show the existence of a multinucleated network of myelin-forming macroglia that projects into neuronal somata and internalizes cellular waste. We demonstrate that AQP4-immunoreactive glial profiles are contained within the myelin sheath and swell in the vicinity of forming waste-canals. Based on these observations we postulate that this process may contribute to an AQP4-medicated bulk flow toward the waste-canals explaining the observed accumulation of lipofuscin in and around waste-canals. We demonstrate that this glial-canal system shows hypertrophic swelling in degenerating brain tissue. These pathologies coincide with the loss of cytoplasmic content and structural integrity of associated neurons whose somata contain myelin-derived waste canals. We demonstrate striking similarities of myelinated glial cells in rodent and human hippocampus. Sequencing the spider genome, we show highly conserved arachnid homologues to human proteins that have been implicated with neurodegeneration. These findings demonstrate the potential of this arachnid model system to study the cellular and functional basis of waste removal and neurodegeneration.

### Proposed functional significance of myelin and AQP4 immunoreactive ACs, GACs and NACs

Our findings show the abundance of lipofuscin in metabolically active neurons highlighting the importance of controlled waste removal to prevent obstruction of neurons or surrounding interstitial spaces. We postulate that the waste-canal system described here consists of two cell types that contribute to the removal of cellular waste from neurons: (*i*) sODG, that form the structural foundation to internalize and sequester lipofuscin, and (*ii*) AQP4 expressing tanycyte-like cells that reside in the lining of a previously described dorsal tubular system. These cells give rise to long tubular processes that pull into the CNS [6]. We propose that these processes are the large diameter ACs described here, and that ACs branch and project into the myelin sheath thus constituting the GACs shown here. As glial membranes and lobes detach on the neuron-facing side of sODG it is plausible that GACs translocate into the neuronal cytoplasm in form of the NACs shown here. We postulate that this translocation triggers the activation of the AQP4 aqua-channels which leads to the swelling of NACs. The proposed interconnected nature of ACs, GACs and NACs is supported by the observed spongiform swelling of all three components in degenerating brain tissue. Based on this model the localized activation of NACs would create a convective flow toward the waste-canals explaining the accumulation of lipofuscin around and within waste-canals shown here. We postulate that the hypertrophic abnormalities may be due to waste obstructions within this waste-canal system.

### Proposed formation of waste-canals

As discussed previously in more detail [6] we postulate that MABs together with continuously forming glial membranes may be the foundation for the continuous supply of waste-canals throughout the lifespan of these animals. We postulate that myelin-forming membranes that initially appear circular around forming glial lobes [6] are enzymatically cleaved and anchored to microtubules via microtubule associated proteins, the identity of these proteins is currently unknown. This process would provide a controlled mechanism by which myelin membranes can be partially detached from the main myelin lobe and form the waste-canals described here. The (*i*) dynamic instability of heterodimeric microtubules, and (*ii*) the interaction of microtubules with a large body of associated proteins make this cytoskeletal filament uniquely suitable for the dynamic attachment/detachment of one or more myelin membranes from their microtubule anchors [23, 24]. It is feasible that such processes may be governed by biochemical processes associated with increased waste accumulation, such as localized lysosome-induced changes in cytoplasmic pH that triggers enzymatic activation/inactivation [25, 26]. To test these postulations, the assembly of the spider genome together with altered gene and protein expression in healthy and degenerating spider brain will enable us to investigate altered expression/phosphorylation of microtubule associated proteins.

It is of particular interest that AD is associated with the accumulation of hyperphosphorylated tau protein [27]. In this context it is imperative to investigate the anatomical foundation of microtubules and their role in the structural composition of myelinated ependymal tanycytes.

### The importance of invertebrate model systems in neuroscience

We have utilized giant spider neurons that show the existence of myelin-derived waste-canals with impressive clarity due to the large diameters of both neurons and associated structures. Guided by the discovery of this waste-internalizing glial-canal system we have provided robust evidence that this vitally important process of waste removal from neurons may be highly conserved in both rodent and human brain[4–6]. We argue that our studies are of critical importance as they provide the structural foundation that informs the proposed testable functional hypothesis.

A similar model system with giant neurons in the longfin inshore squid *Doryteuthis pealeii* was utilized by Hodgkin and Huxley to elucidate the underlying mechanisms that generate and conduct action potentials [28–31]. Inspired by these discoveries it was later demonstrated that the same functional mechanism are highly conserved in amphibian [32] and mammalian systems [33]. Similarly, our understanding regarding the underlying mechanisms of learning and memory were significantly advanced by studies of the gill-withdrawal reflex in the sea slug *Aplysia californica* [34]. Despite evolutionary adaptations in higher developed organisms, the initial discovery of these systems in invertebrate model systems has significantly advanced our knowledge of such systems [35].

### The role of myelin in the nervous system

One of the most significant findings of this study is the waste-internalizing role of myelin shown here. Based on the current understanding the functional significance of myelin is the insulation of neuronal axons to facilitate rapid signal transduction. The current dogma is that myelin is formed by mononucleated oligodendroglia whose processes wrap around neuronal axons [36–39]. The observation of multinucleated ependymal tanycytes that give rise to vast numbers of varicose myelinated processes that project parallel to unmyelinated axons of hippocampal pyramidal cells (see also Figure 6 in [6]) is inconsistent with our current understanding of myelination. We have unambiguously identified axonal processes based on their physical association with the neuronal somata in the well-described hippocampal formation. This inconsistency highlights the importance of re-investigating the ultrastructure, including immunoelectron microscopy of the nervous system. Currently varicose formations along myelinated cell profiles are considered fixation artifacts [37, 40]. Immunolabeling for AQP4 at the EM level will be invaluable to investigate this topic. To this end sequencing of the spider genome will allow us to generate relevant antibodies that are suitable for electron microscopic fixation protocols.

### The *C. salei* genome shows consistency with other arachnid species

Regarding the reported draft mitochondrial genome in *C. salei*, the orientation of the reported elements is nearly identical to that of *Heliophanus lineiventris* (Salticidae) [41], although the *C. salei* mitochondria lack tRNA-A. However, these findings are not unexpected since Araneae are known to contain mitochondrial tRNA gene losses and reductions in tRNA structure [42]. The genome size of C. salei falls within the range of sequenced spider genomes [43] and is comparable to the golden orb-weaver, *Nephila clavipes* [43, 44].

## Abbreviations

Ab: Amyloid b
ACs: Aqua-canals
AD: Alzheimer disease
AQP4: Aquaporin 4
GACs: Glial aqua channel
GM: Geometric means
MABs: Microtubule-associated breaking points
NACs: Neuronal aqua channel
PBS: Phosphate buffered saline
sODG: Spider Oligodendroglia

## ACKNOWLEDGEMENTS

The authors would like to thank Drs. Christopher Francklyn, Douglas Taatjes, Mark Nelson, and Mark Lubkowitz, for support and helpful feedback on this work. We thank Ann and Dr. David MacLaughlin for their generous donation to fund the Zeiss confocal microscope used here. We thank the Vermont Center for Cardiovascular and Brain Health, the Vermont Integrated Genomics Resource and Vermont Advanced Computing Center for scientific support. Electron Microscopic work was performed at the Microscopy Imaging Center at the University of Vermont (RRID: SCR_018821). We thank Nicole Bouffard, and Brad Vietje for expert feedback and technical support. Research reported in this publication was supported by an Institutional Development Award (IDeA) from the National Institute of General Medical Sciences of the National Institutes of Health under grant number P20GM103449. Its contents are solely the responsibility of the authors and do not necessarily represent the official views of NIGMS or NIH.

## AUTHOR CONTRIBUTIONS

Conceptualization: RFF

Funding acquisition: RFF, ALW, JAD

Methodology: RFF, ALW, HED, EEC, JAL, EEC

Project administration: RFF, EEC

Supervision: RFF, EEC

Investigation: RFF, HED, SMBG, MNC, TAD, IRD, MDD, ZFF, KMH, AMH, ICJ, LGK, LKK, KJL, LAK, JAL, JSM, SCM, BAM, OAM, JAM, CMP, OVP, ESSP, CJR, KCS, HAS, JES, MEW, AEW, ALW, ALW, EEC

Visualization: RFF, ALW, HED, SMBG, IRD, KMH, LKK, LAK, BAM, CMP, KCS, HAS, MEW, AEW, EEC

Writing first draft: RFF

Editing: SMBG, MNC, TAD, MDD, JAD, HED, ZFF, ICJ, SCM, OAM, JAM, OVP, ESSP, CJR, JES, EEC

---

**Figure S1.**
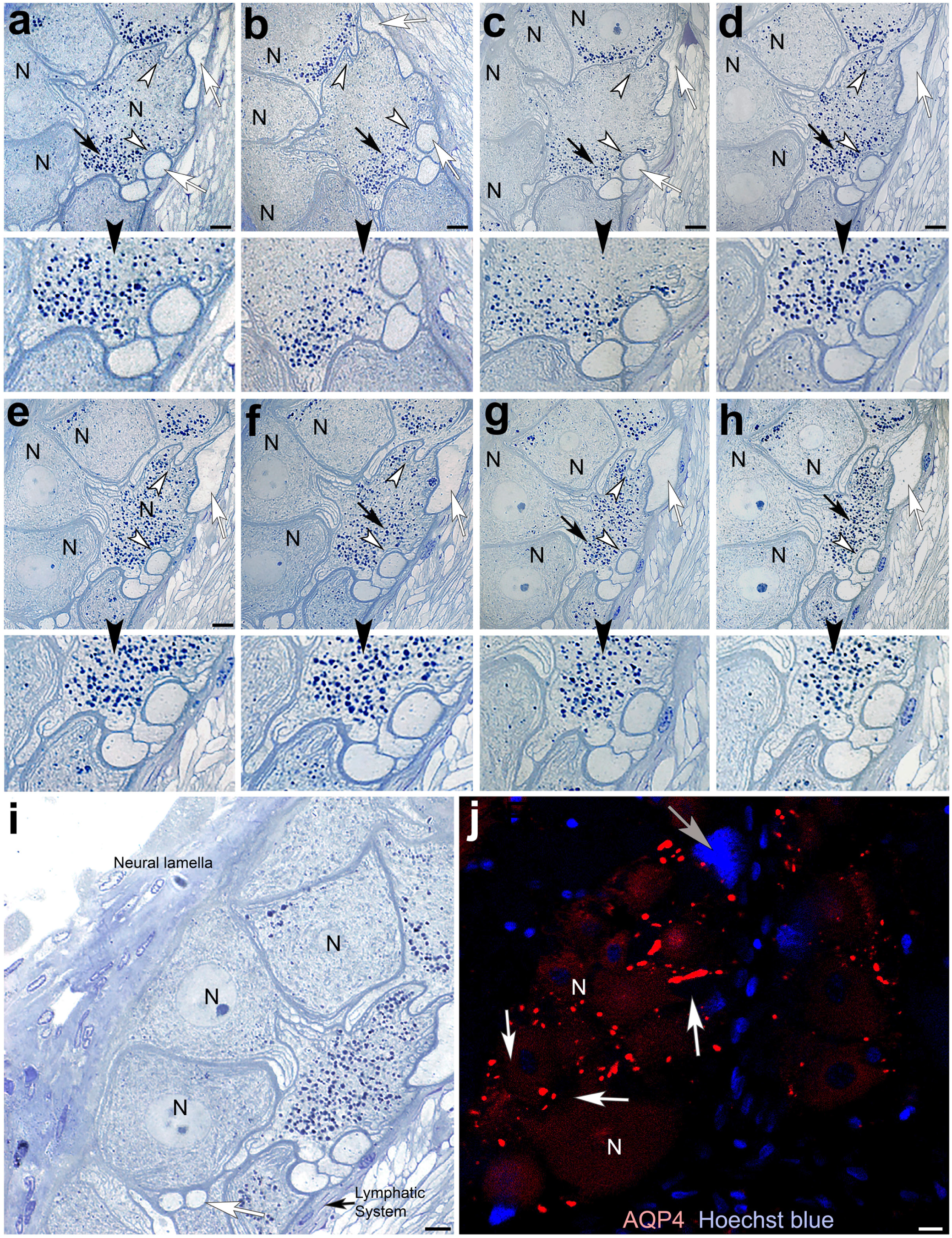
Serial sections of toluidine blue stained neurons in healthy leg ganglia of *Cupiennius salei*. **(a-i)** Neuronal somata (*N*) show large accumulations of granular lipofuscin particles (*black arrows*) in the vicinity of myelin derived waste-canals (*white arrowheads*). Higher zoom shown in the insets below each picture (*black arrowheads*) reveals the internalization of waste particles by glial canals. **(j)** Aquaporin-4 immunolabeling of neurons in leg ganglia shows punctate immunolabeling of canal structures around neurons (*white arrows*). Scale bars: 10 μm.

**Figure S2.**
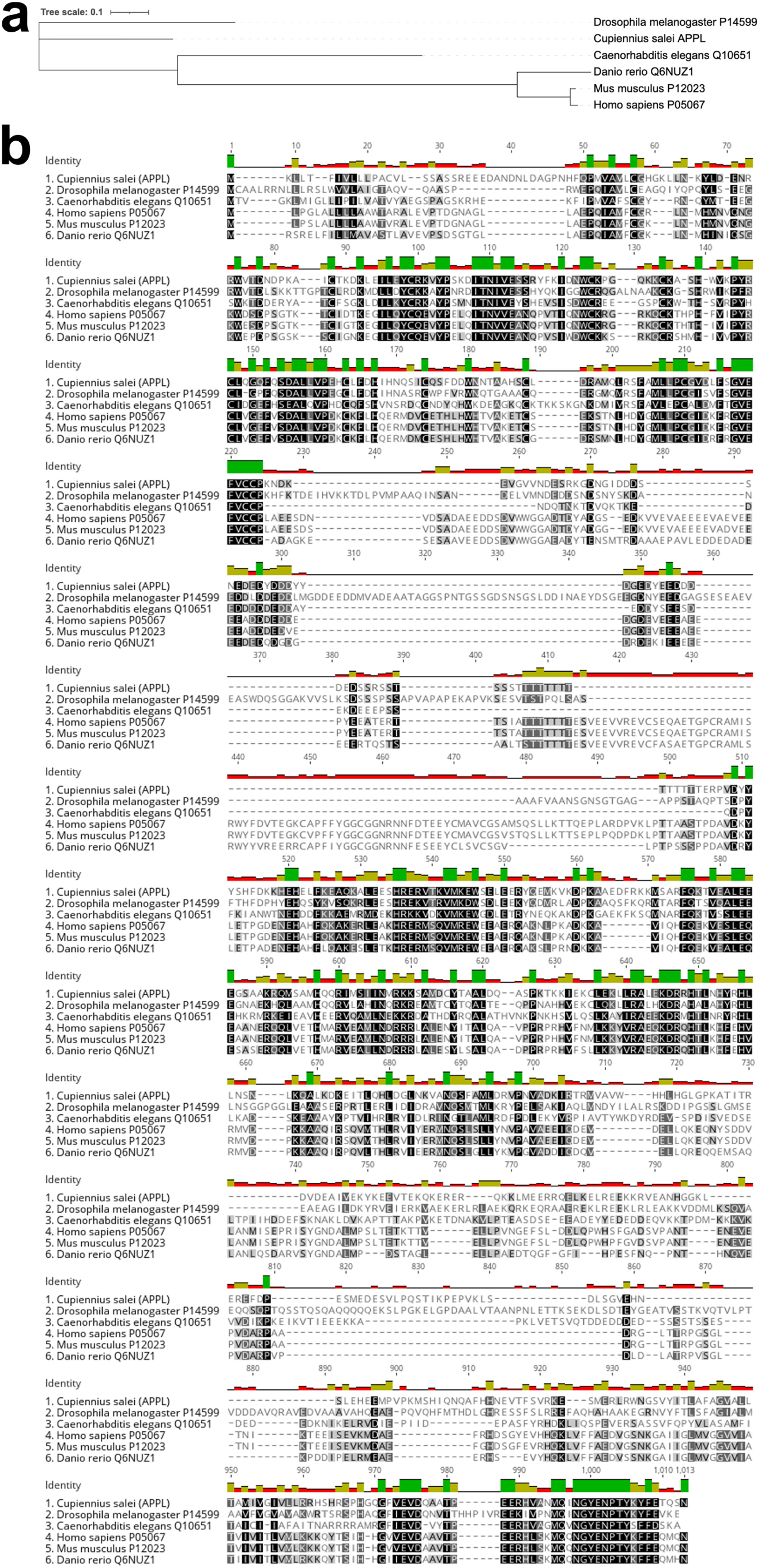
Maximum likelihood phylogenetic tree and amino acid multiple sequence alignment for Amyloid-β (APPL/APP/APPA/APL1).

**Figure S3.**
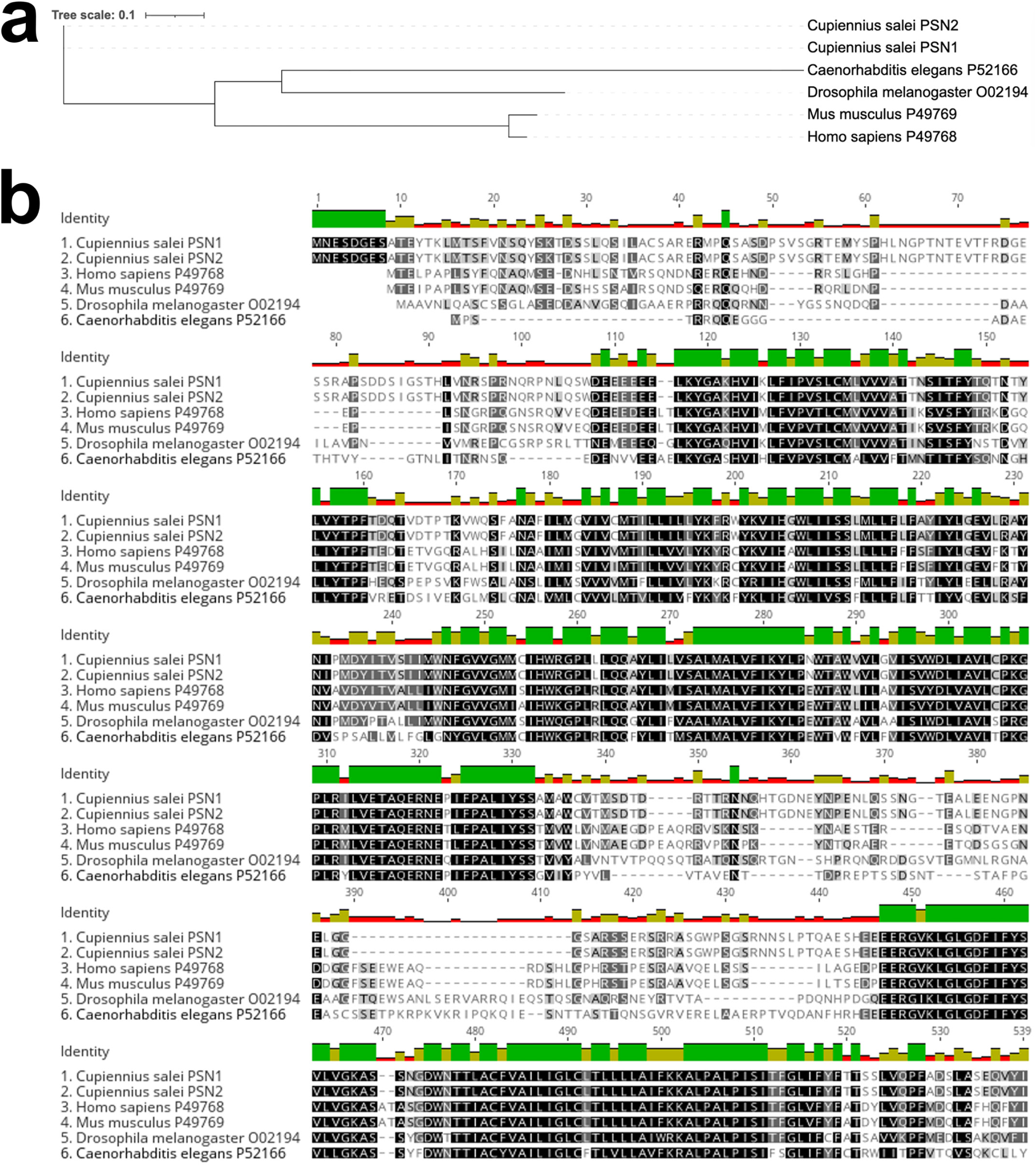
Maximum likelihood phylogenetic tree and amino acid multiple sequence alignment for γ-secretase complex: presenilin (PSN/PSEN1/SEL-12).

## REFERENCES

1. Crous-Bou M, Minguillon C, Gramunt N, Molinuevo JL. Alzheimer’s disease prevention: from risk factors to early intervention. Alzheimers Res Ther. 2017;9(1):71. Epub 20170912. doi: 10.1186/s13195-017-0297-z. PubMed PMID: 28899416; PubMed Central PMCID: PMC5596480.

2. Hardy JA, Higgins GA. Alzheimer’s disease: the amyloid cascade hypothesis. Science. 1992;256(5054):184–5. doi: 10.1126/science.1566067. PubMed PMID: 1566067.

3. Selkoe DJ, Hardy J. The amyloid hypothesis of Alzheimer’s disease at 25 years. EMBO Mol Med. 2016;8(6):595–608. Epub 20160601. doi: 10.15252/emmm.201606210. PubMed PMID: 27025652; PubMed Central PMCID: PMC4888851.

4. Fabian-Fine R, Roman AG, Weaver AL. Alzheimer Disease: The proposed role of tanycytes in the formation of tau tangles and amyloid beta plaques in human brain. bioRxiv. 2025:2025.04.08.647836. doi: 10.1101/2025.04.08.647836.

5. Fabian-Fine R, Roman AG, Winters MJ, Lathram KJ, Bennett CH, Kipingi LK, et al. Uncovering the invisible giant: Amyloid ss plaques and their proposed association with waste removal in Alzheimer-affected human hippocampus. bioRxiv. 2025. Epub 20250606. doi: 10.1101/2025.06.01.657219. PubMed PMID: 40501787; PubMed Central PMCID: PMC12157629.

6. Fabian-Fine R, Weaver AL, Roman AG, Winters MJ, DeWitt JC. Myelinated Glial Cells: Their Proposed Role in Waste Clearance and Neurodegeneration in Arachnid and Human Brain. J Comp Neurol. 2024;532(11):e70000. doi: 10.1002/cne.70000. PubMed PMID: 39610046; PubMed Central PMCID: PMC11605019.

7. Nedergaard M. Neuroscience. Garbage truck of the brain. Science. 2013;340(6140):1529–30. doi: 10.1126/science.1240514. PubMed PMID: 23812703.

8. Iliff JJ, Nedergaard M. Is there a cerebral lymphatic system? Stroke. 2013;44(6 Suppl 1):S93–S5. doi: 10.1161/STROKEAHA.112.678698. PubMed PMID: 23709744.

9. Iliff JJ, Wang M, Liao Y, Plogg BA, Peng W, Gundersen GA, et al. A paravascular pathway facilitates CSF flow through the brain parenchyma and the clearance of interstitial solutes, including amyloid β. Science Translational Medicine. 2012;4(147):147ra11. Epub 2012/08/17. doi: 10.1126/scitranslmed.3003748. PubMed PMID: 22896675; PubMed Central PMCID: PMC3551275.

10. Ghanizada H, Nedergaard M. The glymphatic system. Handb Clin Neurol. 2025;209:161–70. doi: 10.1016/B978-0-443-19104-6.00006-1. PubMed PMID: 40122623.

11. Kumar D, Sharma A, Sharma L. A Comprehensive Review of Alzheimer’s Association with Related Proteins: Pathological Role and Therapeutic Significance. Curr Neuropharmacol. 2020;18(8):674–95. doi: 10.2174/1570159X18666200203101828. PubMed PMID: 32172687; PubMed Central PMCID: PMC7536827.

12. Camacho C, Coulouris G, Avagyan V, Ma N, Papadopoulos J, Bealer K, et al. BLAST+: architecture and applications. BMC Bioinformatics. 2009;10:421. Epub 20091215. doi: 10.1186/1471-2105-10-421. PubMed PMID: 20003500; PubMed Central PMCID: PMC2803857.

13. UniProt C. UniProt: a hub for protein information. Nucleic Acids Res. 2015;43(Database issue):D204–12. Epub 20141027. doi: 10.1093/nar/gku989. PubMed PMID: 25348405; PubMed Central PMCID: PMC4384041.

14. Alliance of Genome Resources C. Updates to the Alliance of Genome Resources Central Infrastructure Alliance of Genome Resources Consortium. bioRxiv. 2023. Epub 20231122. doi: 10.1101/2023.11.20.567935. PubMed PMID: 38045425; PubMed Central PMCID: PMC10690154.

15. Katoh K, Standley DM. MAFFT multiple sequence alignment software version 7: improvements in performance and usability. Mol Biol Evol. 2013;30(4):772–80. Epub 20130116. doi: 10.1093/molbev/mst010. PubMed PMID: 23329690; PubMed Central PMCID: PMC3603318.

16. Steenwyk JL, Buida TJ, 3rd, Li Y, Shen XX, Rokas A. ClipKIT: A multiple sequence alignment trimming software for accurate phylogenomic inference. PLoS Biol. 2020;18(12):e3001007. Epub 20201202. doi: 10.1371/journal.pbio.3001007. PubMed PMID: 33264284; PubMed Central PMCID: PMC7735675.

17. Kalyaanamoorthy S, Minh BQ, Wong TKF, von Haeseler A, Jermiin LS. ModelFinder: fast model selection for accurate phylogenetic estimates. Nat Methods. 2017;14(6):587–9. Epub 20170508. doi: 10.1038/nmeth.4285. PubMed PMID: 28481363; PubMed Central PMCID: PMC5453245.

18. Minh BQ, Schmidt HA, Chernomor O, Schrempf D, Woodhams MD, von Haeseler A, et al. IQ-TREE 2: New Models and Efficient Methods for Phylogenetic Inference in the Genomic Era. Mol Biol Evol. 2020;37(5):1530–4. doi: 10.1093/molbev/msaa015. PubMed PMID: 32011700; PubMed Central PMCID: PMC7182206.

19. Zhou L, Feng T, Xu S, Gao F, Lam TT, Wang Q, et al. ggmsa: a visual exploration tool for multiple sequence alignment and associated data. Brief Bioinform. 2022;23(4). doi: 10.1093/bib/bbac222. PubMed PMID: 35671504.

20. Wang LG, Lam TT, Xu S, Dai Z, Zhou L, Feng T, et al. Treeio: An R Package for Phylogenetic Tree Input and Output with Richly Annotated and Associated Data. Mol Biol Evol. 2020;37(2):599–603. doi: 10.1093/molbev/msz240. PubMed PMID: 31633786; PubMed Central PMCID: PMC6993851.

21. Bankhead P, Loughrey MB, Fernandez JA, Dombrowski Y, McArt DG, Dunne PD, et al. QuPath: Open source software for digital pathology image analysis. Sci Rep. 2017;7(1):16878. Epub 20171204. doi: 10.1038/s41598-017-17204-5. PubMed PMID: 29203879; PubMed Central PMCID: PMC5715110.

22. Schindelin J, Arganda-Carreras I, Frise E, Kaynig V, Longair M, Pietzsch T, et al. Fiji: an open-source platform for biological-image analysis. Nature Methods. 2012;9(7):676–82. doi: 10.1038/nmeth.2019.

23. Billger MA, Bhattacharjee G, Williams RC, Jr. Dynamic instability of microtubules assembled from microtubule-associated protein-free tubulin: neither variability of growth and shortening rates nor “rescue” requires microtubule-associated proteins. Biochemistry. 1996;35(42):13656–63. doi: 10.1021/bi9616965. PubMed PMID: 8885845.

24. Goodson HV, Jonasson EM. Microtubules and Microtubule-Associated Proteins. Cold Spring Harb Perspect Biol. 2018;10(6). Epub 20180601. doi: 10.1101/cshperspect.a022608. PubMed PMID: 29858272; PubMed Central PMCID: PMC5983186.

25. Zhang J, Zeng W, Han Y, Lee WR, Liou J, Jiang Y. Lysosomal LAMP proteins regulate lysosomal pH by direct inhibition of the TMEM175 channel. Mol Cell. 2023;83(14):2524–39 e7. Epub 20230629. doi: 10.1016/j.molcel.2023.06.004. PubMed PMID: 37390818; PubMed Central PMCID: PMC10528928.

26. Quick JD, Silva C, Wong JH, Lim KL, Reynolds R, Barron AM, et al. Lysosomal acidification dysfunction in microglia: an emerging pathogenic mechanism of neuroinflammation and neurodegeneration. Journal of neuroinflammation. 2023;20(1):185.

27. Iqbal K, Liu F, Gong CX, Grundke-Iqbal I. Tau in Alzheimer disease and related tauopathies. Curr Alzheimer Res. 2010;7(8):656–64. doi: 10.2174/156720510793611592. PubMed PMID: 20678074; PubMed Central PMCID: PMC3090074.

28. Hodgkin AL, Huxley AF. A quantitative description of membrane current and its application to conduction and excitation in nerve. J Physiol. 1952;117(4):500–44. doi: 10.1113/jphysiol.1952.sp004764. PubMed PMID: 12991237; PubMed Central PMCID: PMC1392413.

29. Hodgkin AL, Huxley AF. Currents carried by sodium and potassium ions through the membrane of the giant axon of Loligo. J Physiol. 1952;116(4):449–72. doi: 10.1113/jphysiol.1952.sp004717. PubMed PMID: 14946713; PubMed Central PMCID: PMC1392213.

30. Hodgkin AL, Huxley AF. The components of membrane conductance in the giant axon of Loligo. J Physiol. 1952;116(4):473–96. doi: 10.1113/jphysiol.1952.sp004718. PubMed PMID: 14946714; PubMed Central PMCID: PMC1392209.

31. Hodgkin AL, Huxley AF. The dual effect of membrane potential on sodium conductance in the giant axon of Loligo. J Physiol. 1952;116(4):497–506. doi: 10.1113/jphysiol.1952.sp004719. PubMed PMID: 14946715; PubMed Central PMCID: PMC1392212.

32. Hille B. Pharmacological modifications of the sodium channels of frog nerve. J Gen Physiol. 1968;51(2):199–219. doi: 10.1085/jgp.51.2.199. PubMed PMID: 5641635; PubMed Central PMCID: PMC2201123.

33. Beneski DA, Catterall WA. Covalent labeling of protein components of the sodium channel with a photoactivable derivative of scorpion toxin. Proc Natl Acad Sci U S A. 1980;77(1):639–43. doi: 10.1073/pnas.77.1.639. PubMed PMID: 6928649; PubMed Central PMCID: PMC348330.

34. Bailey CH, Kandel ER. Synaptic remodeling, synaptic growth and the storage of long-term memory in Aplysia. Prog Brain Res. 2008;169:179–98. doi: 10.1016/S0079-6123(07)00010-6. PubMed PMID: 18394474.

35. Orvis J, Albertin CB, Shrestha P, Chen S, Zheng M, Rodriguez CJ, et al. The evolution of synaptic and cognitive capacity: Insights from the nervous system transcriptome of Aplysia. Proc Natl Acad Sci U S A. 2022;119(28):e2122301119. Epub 20220708. doi: 10.1073/pnas.2122301119. PubMed PMID: 35867761; PubMed Central PMCID: PMC9282427.

36. Boullerne AI. The history of myelin. Experimental Neurology. 2016;283(Part B):431–45. doi: 10.1016/j.expneurol.2016.06.005. PubMed PMID: S0014488616301704.

37. Möbius W, Nave K-A, Werner HB. Electron microscopy of myelin: Structure preservation by high-pressure freezing. Brain Research. 2016;1641(Part A):92–100. doi: 10.1016/j.brainres.2016.02.027. PubMed PMID: 2016-11609-001.

38. Simons M, Nave K-A. Oligodendrocytes: Myelination and Axonal Support. Cold Spring Harbor perspectives in biology. 2015;8(1):a020479. doi: 10.1101/cshperspect.a020479. PubMed PMID: 26101081.

39. Stadelmann C, Timmler S, Barrantes-Freer A, Simons M. Myelin in the central nervous system: structure, function, and pathology. Physiological reviews. 2019;99(3):1381–431.

40. Möbius W, Cooper B, Kaufmann WA, Imig C, Ruhwedel T, Snaidero N, et al. Electron microscopy of the mouse central nervous system. Methods in cell biology. 96: Elsevier; 2010. p. 475–512.

41. Li M, Chen W-T, Zhang Q-L, Liu M, Xing C-W, Cao Y, et al. Mitochondrial phylogenomics provides insights into the phylogeny and evolution of spiders (Arthropoda: Araneae). Zoological Research. 2022;43(4):566.

42. Ozerova I, Fallmann J, Morl M, Bernt M, Prohaska SJ, Stadler PF. Aberrant Mitochondrial tRNA Genes Appear Frequently in Animal Evolution. Genome Biol Evol. 2024;16(11). doi: 10.1093/gbe/evae232. PubMed PMID: 39437314; PubMed Central PMCID: PMC11571959.

43. Sharma PP. The Impact of Whole Genome Duplication on the Evolution of the Arachnids. Integr Comp Biol. 2023;63(3):825–42. doi: 10.1093/icb/icad050. PubMed PMID: 37263789.

44. Babb PL, Lahens NF, Correa-Garhwal SM, Nicholson DN, Kim EJ, Hogenesch JB, et al. The Nephila clavipes genome highlights the diversity of spider silk genes and their complex expression. Nature Genetics. 2017;49(6):895–903.

